# The C-terminal domains SnRK2-box and ABA-box have a role in sugarcane SnRK2s auto-activation and activity

**DOI:** 10.1101/589903

**Authors:** Germanna Righetto, Dev Sriranganadane, Levon Halabelian, Carla G. Chiodi, Jonathan M. Elkins, Katlin B. Massirer, Opher Gileadi, Marcelo Menossi, Rafael M. Couñago

## Abstract

Resistance to drought stress is fundamental to plant survival and development. Abscisic acid (ABA) is one of the major hormones involved in different types of abiotic and biotic stress responses. ABA intracellular signaling has been extensively explored in *Arabidopsis thaliana* and occurs via a phosphorylation cascade mediated by three related protein kinases, denominated SnRK2s (SNF1-related protein kinases). However, the role of ABA signaling and the biochemistry of SnRK2 in crop plants remains underexplored. Considering the importance of the ABA hormone in abiotic stress tolerance, here we investigated the regulatory mechanism of sugarcane SnRK2s - known as SAPKs (Stress/ABA-activated Protein Kinases). The crystal structure of ScSAPK10 revealed the characteristic SnRK2 family architecture, in which the regulatory SnRK2-box interacts with the kinase domain αC helix. To study sugarcane SnRK2 regulation, we produced a series of mutants for the protein regulatory domains SnRK2-box and ABA-box. Surprisingly, mutations in the SnRK2-box did not drastically affect sugarcane SnRK2 activity, in contrast to previous observations for the homologous proteins in Arabidopsis. Also, we found that the ABA-box might have a role in SnRK2 activation in the absence of PP2C phosphatase. Taken together, our results demonstrate that both C-terminal domains of sugarcane SnRK2 proteins play a fundamental role in protein activation and activity.

## INTRODUCTION

The phytohormone abscisic acid (ABA) is a central regulator of plant responses to abiotic stress. ABA triggers protective plant responses leading to stomatal closure, seed dormancy, inhibition of growth, and germination (Fujii et al., 2007; Fujii and Zhu, 2009; Mustilli, 2002; Yoshida et al., 2002, 2006). ABA’s signaling role is carried out by a protein phosphorylation cascade that depends on the interplay between the activities of SnRK2 kinases and protein phosphatase 2C (PP2C) (Fujii et al., 2009; Umezawa et al., 2010).

In dicotyledons, members of the SnRK2 sub-family of serine-threonine kinases (SnRK2.2/2.3/2.6 in Arabidopsis) act as positive regulators of ABA signaling and activate downstream stress-responsive genes and transcription factors (Fujii et al., 2009; Fujii and Zhu, 2009; Fujita et al., 2009; Nakashima et al., 2009). Counterpart kinases in monocots are known as SAPK8/9/10 (Stress/ABA-activated Protein Kinases) (Belin et al., 2006; Kobayashi et al., 2004; Yoshida et al., 2006). ABA-responsive kinases from both mono- and dicotyledons are expected to have a conserved modular architecture and to be involved in environmental sensing and stress response (Kulik et al., 2011).

The C-terminal SnRK2-box is essential for kinase activation by hyperosmotic stress and displays high sequence conservation among members of the SnRK2 subfamily (Kobayashi et al., 2004; Yoshida et al., 2006). The crystallographic structures of Arabidopsis SnRK2.3 and 2.6 have shown that the SnRK2-box folds into a helix and packs against the catalytically important αC helix within the protein kinase domain (Ng et al., 2011; Yunta et al., 2011). Mutational studies have demonstrated that the interaction between these two helices is crucial for kinase autoactivation and subsequent phosphorylation of the transcription factor ABF2 (Ng et al., 2011).

SnRK2/SAPK activity is modulated by direct interaction with PP2C phosphatases, which, in turn, depends on intracellular ABA levels. In presence of the hormone, the phosphatase activity is impaired by the interaction with the complex formed by ABA and PYL/PYR/RCAR receptors (Ma et al., 2009; Melcher et al., 2009; Miyazono et al., 2009; Nakashima et al., 2009; Park et al., 2009; Umezawa et al., 2009; Yin et al., 2009). The complex blocks PP2C substrate entry and prevents SnRK2 inactivation by dephosphorylation. In the absence of ABA, PP2C is released from the complex with PYL/PYR/RCAR receptors and can interact with the SnRK2 kinases, leading to kinase dephosphorylation and repression of ABA-response. The interaction between kinase and phosphatase is mediated by another C-terminal motif, known as ABA-box, only preserved in the ABA-responsive members of the SnRK2 subfamily (Soon et al., 2012; Umezawa et al., 2009; Vlad et al., 2009).

Despite extensive characterization in Arabidopsis, the protein structure and biochemical regulation of ABA-responsive SnRK2s from crop plants remain poorly explored. SnRK2 subfamily members have been identified in several crop plants, such as rice, maize and cotton (Huai et al., 2008; Kobayashi et al., 2004; Liu et al., 2017). Just like their counterparts from Arabidopsis, these proteins have been shown to mediate plant responses to abiotic stress and ABA. In *Saccharum officinarum* L. (So) sugarcane, a recent study identified ten SnRK2 subfamily members, three of which (SoSAPK8/9/10) have the characteristic ABA-box in their C-terminus and, accordingly, are responsive to ABA (Li et al., 2017). Despite these studies, currently, there is no structural information on SnRK2 subfamily members from crop plants. Moreover, the role of the regulatory domains SnRK2-box and ABA-box in protein activity and activation remain unclear for sugarcane and other crop plants.

In this study, we report the crystal structure of SAPK10 from the crop plant sugarcane (*Saccharum* ssp. hybrids). We also investigated how SnRK2- and ABA-boxes modulate the activity of SAPK8/9/10. These analyses confirmed that, overall, the SnRK2-box within sugarcane SAPKs preserves its role in protein activity, albeit to a lesser extent when compared to the Arabidopsis proteins. Finally, we identified several auto-phosphorylated sites within SAPK kinase surface that might have a role in their interaction with PP2C and/or downstream partners.

## MATERIAL AND METHODS

### Gene identification and bioinformatics analyses

The sequences of *ScSAPK8, ScSAPK9* and *ScSAPK10* were identified using the Sugarcane Expressed Sequence Tag (SUCEST) database and the homologous sequences from *Sorghum bicolor* (*SbSAPK8*: Sb01g007120, *SbSAPK9*: Sb08g019700 and *SbSAPK10*: Sb01g014720 and *Arabidopsis thaliana* (*SnRK2.2*: 824214, *SnRK2.3*: 836822, *SnRK2.6*: 829541) as reference (Vettore et al., 2003). The coding sequences of the three sugarcane SAPKs were isolated from the sugarcane leaf cDNA (cultivar SP80-3280) using specific primers (Supplementary Table S1).

For analysis of protein conservation, protein sequences from *Arabidopsis thaliana*, and *Saccharum* spp were aligned using BioEdit and Clustal Omega (Hall, 1999; Sievers and Higgins, 2014). The sequence similarities, as well as the secondary structure elements, were further analyzed using the ESPript 3.0 program (Robert and Gouet, 2014). The analysis of protein domains was performed using PFAM and SMART databases (Finn et al., 2016; Schultz et al., 1998).

### *ScSAPKs* cloning and recombinant protein expression in *Escherichia coli*

The full-length sequences of *ScSAPK8/9/10* were cloned into pNIC28-Bsa4 using the ligase-independent cloning (LIC) method (Savitsky et al., 2010). For large scale protein expression, the constructs were transformed into *E. coli* strain BL21(DE3)-R3-pRARE2 (Savitsky et al., 2010) and grown in 20 mL of LB medium with kanamycin (50 mg/mL) and incubated at 37°C. After overnight growth, the bacterial culture was inoculated into 1.5 L of Terrific Broth medium with kanamycin (50 mg/mL), which was incubated at 37°C with shaking until an OD_600_ of 1.5. The culture was cooled to 18°C before the addition of 0.2 mM of IPTG (Isopropyl β-D-1-thiogalactopyranoside) for overnight expression. Cells were harvested by centrifugation at 7,500 ×*g* at 4°C and suspended in approximately 20 mL of 2X lysis buffer (100 mM HEPES pH 7.5; 1 M NaCl, 20 mM imidazole, 20% glycerol) with 1 μL per mL protease inhibitor cocktail. Suspended cells were placed on ice and sonicated for 9 min (5 s ON; 10 s OFF; 30% amplitude). Polyethyleneimine (pH 7.5) was added to the lysate at 0.15 % final concentration and the lysate was clarified by centrifugation at 53,000 ×g for 45 min at 4°C. The supernatant was loaded onto an IMAC column (5 mL HisTrap FF Crude) and washed with Binding Buffer (50 mM HEPES pH 7.4, 500 mM NaCl, 5% glycerol, 10 mM imidazole pH 7.4, 0.5 mM tris(2-carboxyethyl)phosphine (TCEP)) and Wash Buffer (50 mM HEPES pH 7.4, 500 mM NaCl, 5% glycerol, 30 mM imidazole pH 7.4, 0.5 mM TCEP). The protein was eluted with 10 mL of Elution Buffer (50 mM HEPES pH 7.4, 500 mM NaCl, 5% glycerol, 300 mM imidazole pH 7.4, 0.5 mM TCEP) in 2 mL fractions. The eluted fractions were combined and incubated with TEV protease during overnight dialysis against GF Buffer (Binding Buffer without imidazole). TEV protease, as well as the cleaved 6xHis-tag, were removed using nickel-affinity chromatography resin. The protein was concentrated to 5 mL with a 30 kDa MWCO spin concentrator and loaded onto a size exclusion HiLoad 16/60 Superdex 200pg (GE) column equilibrated in GF buffer. Fractions of 1.8 mL were collected and verified for protein purity on a 12% SDS-PAGE gel. Purified fractions were combined, concentrated and stored at −80°C.

### ScSAPK10 crystallization, data collection and structure determination

For crystallization experiments, the truncated construct of ScSAPK10 corresponding to amino acids 12 to 320 (ScSAPK10_ΔNterm-ΔABA-box) was cloned and the recombinant protein produced as above. Before setting up crystallization trials, protein aliquots at 24 mg/mL were thawed and centrifuged at 15,000 rpm for 10 min at 4 ° C. Crystallization sitting drops were manually mounted using 1:1 ratio of protein to reservoir solution (1.5M ammonium sulfate; 0.1M bis-tris pH 6.5 and 0.1M sodium chloride– *Index Screen*, Hampton Research). Crystals grew after 2 days at 20 °C and were cryoprotected in reservoir solution supplemented with 30% glycerol before flash-cooling in liquid nitrogen. Diffraction data was collected at the Advanced Photon Source (Chicago, USA) beamline 19ID. The X-ray diffraction data was integrated with XDS (Kabsch, 2010) and scaled using AIMLESS from the CCP4 software suite (Winn et al., 2011). The structure was solved by molecular replacement using Phenix (Adams et al., 2002) and the *Arabidopsis thaliana* SnRK2.6 structure (PDB ID 3ZUT) as the initial model (Yunta et al., 2011). Refinement was performed using REFMAC5 (Murshudov et al., 2011). Coot (Emsley et al., 2010) was used for manual model-building and local refinement. Structure validation was performed using MolProbity (Chen et al., 2010). Structure coordinates have been deposited in the Protein Data Bank (PDB ID 5WAX) (Table 1).

**Table 1:** Data collection and refinement statistics

| **Protein** | **ScSAPK10** |
| --- | --- |
| **PDB ID** | **5WAX** |
| **Data collection** |  |
| X-ray source | APS 19-ID |
| Wavelength (Å) | 0.979200 |
| Space Group | C 2 2 2<sub>1</sub> |
| Cell dimensions (Å) a, b, c. | 75.4, 214.6, 93.8 |
| Cell dimensions (°) α, β, γ. | 90, 90, 90 |
| Molecules/ asymmetric unit | 2 |
| Resolution (Å)* | 46.58 - 2.00 (2.05 - 2.0) |
| Unique reflections* | 51563 (3763) |
| $R_{merge}$ (%)* | 8.3 (98.3) |
| I/σ (I)* | 16.0 (1.6) |
| CC (1/2)* | 0.999 (0.604) |
| Completeness (%)* | 99.6 (99.9) |
| Redundancy* | 5.5 (5.6) |
| **Refinement** |  |
| Resolution (Å)* | 46.58 - 2.00 (2.05 - 2.0) |
| $R_{cryst}/R_{free}$ (%) | 19.83 / 23.7 |
| No. Atoms (protein/ solvent) | 4360 / 387 |
| Mean B-factor (Å<sup>2</sup>) | 29.9 |
| R.m.s.d bond lengths (Å), angles (°) | 0.012, 1.49 |
| **Ramachandran statistics (%)** |  |
| Favored / Allowed / Outliers | 96.3 / 3.7 / 0 |

\*Values in parentheses represent the highest resolution shell

### Site-directed mutagenesis and ScSAPK8 expression for phosphorylation assays

The SnRK2-box and ABA-box mutants were produced by site-directed mutagenesis with specific primers (Supplementary Table S1) using as template the full-length construct of ScSAPK8 cloned in pNIC28-Bsa4. The mutated constructs were confirmed by sequencing and transformed in *E. coli* BL21(DE3)-R3 cells which express rare tRNAs (plasmid pACYC-LIC+) and the *λ*-phosphatase.

All proteins were expressed at the same time using the same protocol described previously. After bacterial culture lysis, the clarified supernatants were loaded in 4 mL of Ni^2+^-sepharose beads (GE Healthcare, Uppsala), washed with Binding Buffer (4 × 4 mL) and Wash Buffer (3 × 4 mL). The proteins were eluted with Elution Buffer (4 × 4 mL), and the imidazole was removed using Sephadex G-25 PD-10 Desalting Columns (GE Healthcare, Uppsala). Protein purity was analyzed by SDS-PAGE gel and protein masses were confirmed by intact mass spectrometry.

### ScSAPK8 WT and mutants autophosphorylation assay

Each protein (diluted in GF buffer to 20 μM final concentration) was incubated with 10 mM MgCl_2_ and 1 mM ATP (Sigma – catalog A7699) at 20°C in a final volume of 200 μL. After every time point (1 hour, 5 hours and overnight), 20 μL of aliquots were removed and the reaction stopped by the addition of 10 mM EDTA. Samples were analyzed by LC-MS. For these assays, the protein concentration was estimated by Bradford (Sigma-Aldrich) and SDS-PAGE analysis (Supplementary Figure S1).

### Kinase activity assay

The enzymatic activity of ScSAPK8 WT and SnRK2-box and ABA-box mutants was measured using a TR-FRET based assay (Cisbio Kinease - catalog #62ST1PEB). Prior to the enzymatic reaction, proteins were diluted in GF buffer supplemented with 10 mM MgCl_2_ to a 20 μM final concentration and incubated overnight at 20° C with or without 1 mM ATP. After 16 hours, the activity of proteins pre-incubated with Mg^2+^/ATP and Mg^2+^ was tested using the peptide STK-1 at 1 μM final concentration. Final assay concentrations were: 50 nM kinase, 2 mM ATP, 10 mM MgCl_2_ and 1 mM DTT. The reaction was allowed to progress for 1 hour at room temperature before the detection step was performed according to the manufacturer’s instructions. FRET signal was acquired using a ClarioStar fluorescence plate reader (BMG Labtech) (excitation/emission wavelengths of 330 and 620/650 nm, respectively). Results reported are from two independent experiments performed in triplicates.

### ScSAPK8 WT phosphosite identification

Purified ScSAPK8 WT and ΔABA-box mutant (20 μM final concentration) were diluted in GF buffer supplemented with 10 mM MgCl_2_ and incubated with 1 mM ATP (Sigma – catalog A7699) overnight at 20°C. The reaction was stopped by adding 10 mM EDTA (final concentration) before samples flash-freezing in liquid nitrogen. Protein intact mass was determined by LC-MS and phosphosites were identified by LC-MS/MS. The sample was buffer-exchanged into 50 mM Ammonium Bicarbonate and treated with 25μL of RapiGest SF (0.2% - Waters Corp. catalog # 186001861) for 15 min at 80°C. Dithiothreitol (DTT - 100 mM stock prepared in 50 mM Ammonium Bicarbonate) was added to a final concentration of 4 mM and the mixture was incubated for 30 min at 60 °C. Iodoacetamide (IAA - 300 mM stock prepared in 50 mM Ammonium Bicarbonate) was added to the mixture at a final concentration of 12 mM. The mixture was protected from light and incubated for 30 min. Trypsin (Promega, Fitchburg, WI, USA - catalog # V511A) prepared in 50 mM Ammonium Bicarbonate was added to the mixture (1:100 mass ratio of trypsin to protein) and incubated for 16 hr at 37°C under agitation. To hydrolyze the RapiGest, Trifluoroacetic acid (TFA - Pierce, Waltham, MA, USA; catalog # 53102) was added and the mixture incubated for 90 min at 37°C. The reaction was centrifuged at 14,000 rpm for 30 min at 6°C and the supernatant transferred to a fresh microcentrifuge tube (Axygen, Union City, CA, USA) for subsequent LC-MSMS analysis.

### Mass spectrometry analysis

For intact mass analysis, samples were analyzed via reverse phase HPLC-ESI-MS in positive ion mode using an Acquity H-class HPLC system coupled to an XEVO G2 Xs Q-ToF mass spectrometer (both from Waters Corp.). A total of 0.5 μL sample (~ 12.5 ng) in mobile phase Solvent A (0.1% formic acid - FA, prepared in water) was applied onto a C4 column (ACQUITY UPLC Protein BEH C4 300 Å, 1.7 μm, 2.1 mm X 100 mm - Waters Corp.) kept at 45 °C. Bound protein was eluted by a gradient of 10-90% Solvent B (0.1% FA in 100% Acetonitrile - ACN) over 4 min. Between each injection, the column was regenerated with 90% Solvent B (for 90 sec) and re-equilibrated to 10% Solvent B (210 sec). Flow rates were 0.5 μL/min for sample application and 0.4 mL/min (wash and elution). For internal calibration, the lockspray properties were: scan time of 0.5 sec; and a mass window of 0.5 Da around Leu-enkephalin (556.2771 Da). The ToF-MS acquisition ranged from 100 to 2,000 Da with a scan time of 1 sec. The cone voltage on the ESI source was fixed at 40 V.

For phosphosite identification, samples were analyzed by reverse phase nanoLC-ESI-MSMS using an Acquity M-class HPLC system coupled to an XEVO G2 Xs Q-ToF (both from Waters Corp.). A total of 2 μL sample in mobile Solvent A was applied onto a Trap column (V/M, Symmetry C18, 100Ǻ, 5 μm, 180 μm × 20 mm) connected to an HSS T3 C18 column (75 μm × 150 mm, 1.8 μm), kept at 45° C. LC was performed at a flow rate of 400 nL/min, and the elution of bound peptides was performed over a 47 min gradient as follows: 0-30.37 min from 7-40% Solvent B; 30.37-32.03 min from 40-85% Solvent B; 32.34-35.34 min at 85% Solvent B; 35.34-37 min from 85-7% Solvent B and 37-47 min at 7% B. The nano-ESI source was set with the following parameters: the capillary voltage was 2.5 kV, the sampling cone and the source offset was set at 30 V, the temperature source was 70 °C, the gas flow and the purge gas were set at 50 and 150 L/h, and the nano gas flow was maintained at 0.5 bar. Data were acquired at 0.5 scan/s, over the mass range of 50-2000 m/z in positive and sensitive mode. The MS data-independent acquisition mode was used with a low energy collision switched off and a high collision energy ramp 15-45 eV in the second function for fragmentation. For mass accuracy, the Glu-Fibrinopeptide (785.84261 Da 2+) was used as lock mass at a concentration of 100 fM (in 40:60 ACN/H_2_O, 0.1% FA) infused at a flow rate of 0.5 μL/min via a lock spray interface and an auxiliary pump. Lock mass scans were acquired every 30 s at a rate of 0.5 scan/s. Lockmass was acquired but not applied on the fly.

### MS data analysis

MS raw data was analyzed using MassLynx v4.1 and processed by MaxEnt 1 (both from Waters Corp.) in order to deconvolute multi-charged combined ion spectra for intact mass analysis. Phosphoproteomic raw data were processed using Protein Lynx Global Server (PLGS, Waters Corp.) against the sugarcane protein database (UniProt release 2017_12). Data processing was performed in two steps. First, PLGS extracted all acquired spectra using the following parameters: lock mass (charge 2= 785.84261 Da/e) window set to 0.4 Da; low energy threshold fixed at 500 counts; elevated energy threshold at 50 counts; chromatographic peak width and MS ToF Resolution were set to automatic. Then, a database search was performed with the following parameters: peptide and fragment tolerance were set to automatic; 2 fragments ion matches per peptide and 5 fragments ion matches per protein were fixed, as well as a minimum of 1 peptide match per protein; one missed cleavage was allowed; trypsin was set as the primary digestion; carbamidomethylation of cysteine was set as a fixed modification, oxidation of methionine and phosphorylation of Ser/Thr/Tyr residues were set as a variable modification.

## RESULTS

### ABA-responsive SnRK2s in sugarcane

Three ABA-responsive SnRK2s were identified within the sugarcane genome (*S. spontaneum* × *S. officinarum* hybrid cultivar) using homologous protein sequences from *Arabidopsis thaliana* and *Sorghum bicolor*. Based on previous studies in monocots and dicots, these were designated ScSAPK8, ScSAPK9 and ScSAPK10 (Boudsocq et al., 2004; Cai et al., 2014; Fujii et al., 2009; Fujita et al., 2009; Kobayashi et al., 2004; LI et al., 2010; Li et al., 2017).

At the amino acid level, sugarcane and *A. thaliana* proteins share high sequence identity (≥ 76 %) (Figure 1). Moreover, ABA-responsive SnRK2s and their sugarcane counterparts display an identical modular architecture, in which the N-terminal kinase domain (KD; ~260 amino acids) is followed by two highly conserved motifs - the SnRK2-box (16 amino acids) and ABA-box (27 amino acids) (Supplementary Figure S2). The function of the N-terminal regions of each protein (about twenty residues) remains to be elucidated.

**Figure 1:**
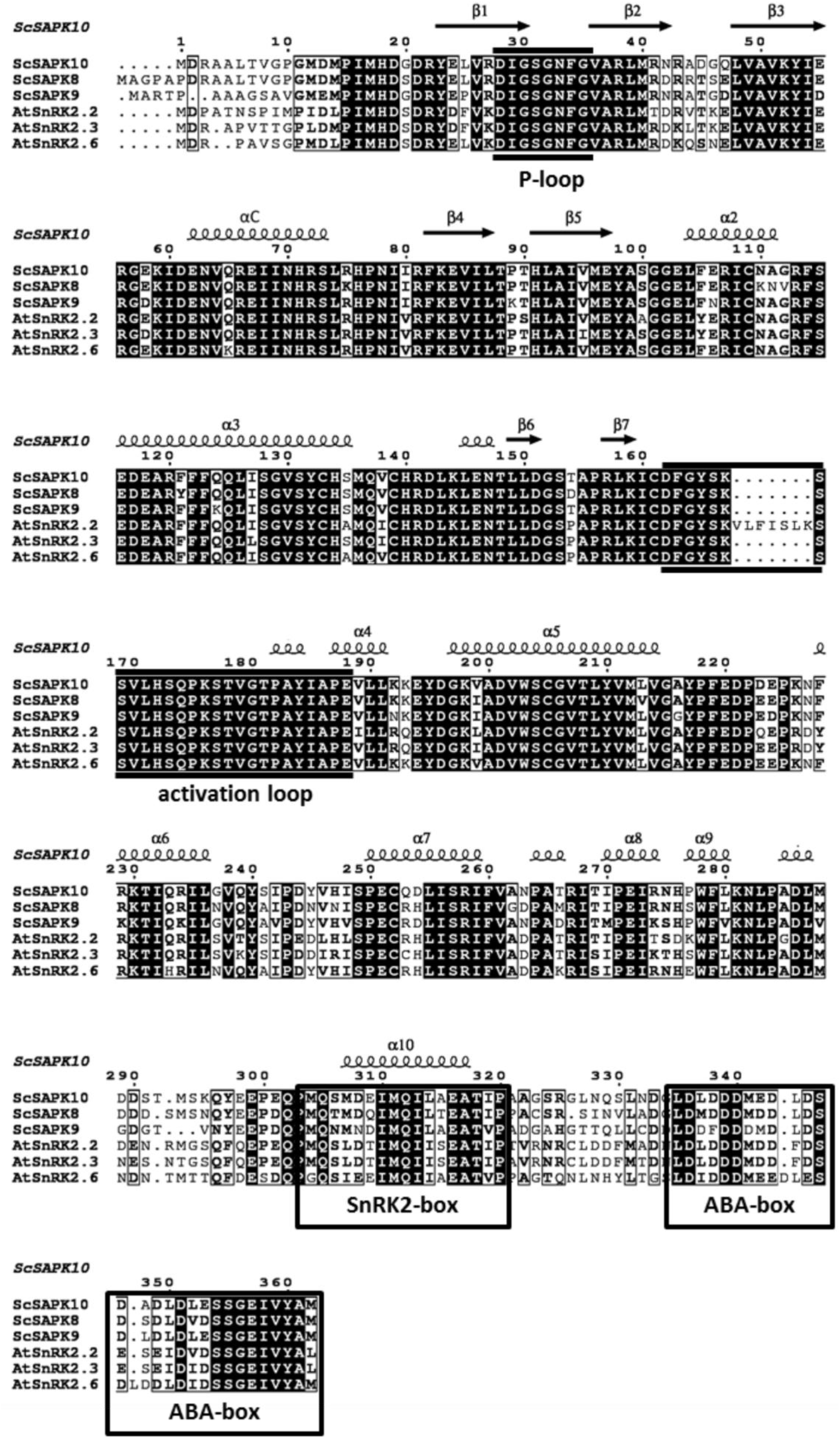
Multiple-sequence alignment of ABA-related SnRK2s shows high similarity and identity between Arabidopsis and sugarcane sequences. Residues colored in black are identical for all the sequences and residues highlighted by a black box share chemical similarity. Sequences corresponding to the P-loop, activation loop and the regulatory domains SnRK2-box and ABA-box are marked based on previous studies (Ng et al., 2011).

### Sugarcane and *A. thaliana* ABA-responsive SnRKs share a conserved kinase fold

To better understand the mechanism of sugarcane ABA-responsive SnRKs, we pursued crystallization of all three ScSAPK proteins. Despite our best efforts, we could not obtain diffraction quality crystals of ScSAPK8 or ScSAPK9. To improve the diffraction quality of initial ScSAPK10 crystals, we used a truncated version of the protein (residues 12 to *320)* in which residues at both N- and C-terminal regions were removed, including the ABA-box. The protein structure was solved at 2.0 Å resolution by molecular replacement using the AtSnRK2.6 structure (PDB ID: 3ZUT) (Yunta et al., 2011) as a model (Table 1).

ScSAPK10 has a canonical kinase fold: a bilobal structure formed by a smaller N-terminal lobe and a larger C-terminal lobe connected by a short hinge region (Figure 2A). The protein N-terminal lobe is composed of five antiparallel β-strands, including the ATP-binding loop (P-loop) between β1 and β2, and the αC helix (Pearce et al., 2010). The C-terminal lobe contains the activation loop and several α-helices. Residues within the ScSAPK10 SnRK2-box (Met304 – Pro320) are folded into an α-helix and packed against αC from the protein kinase domain (Figure 2B). No electron density was observed for residues in the activation loop (residues 165 to 181) or the region of the protein connecting the kinase domain and the SnRK2-box (residues 279 to 294), likely due to flexibility. These regions were omitted from the final model.

**Figure 2:**
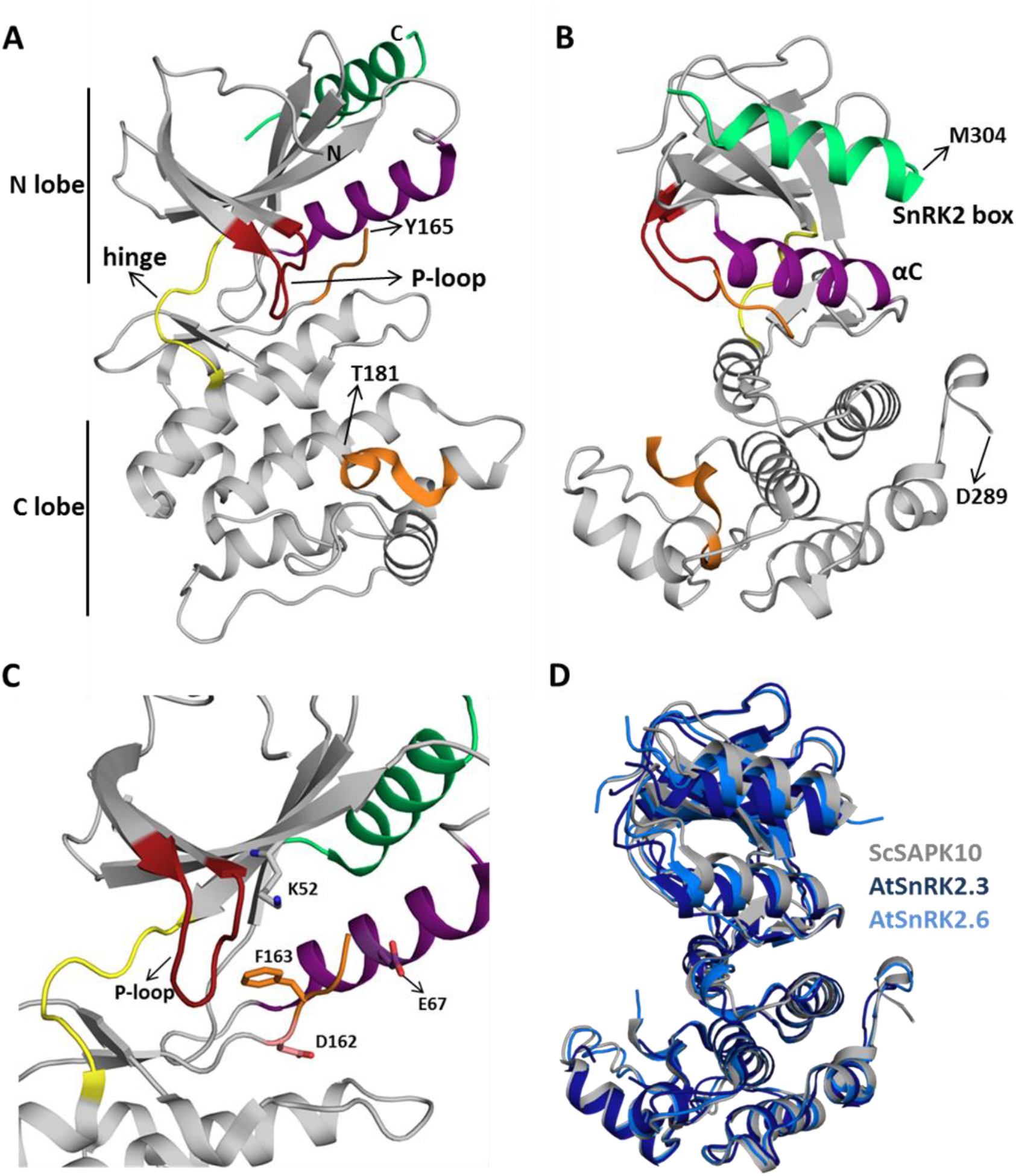
Sugarcane SAPK10 has a canonical kinase fold and shows conserved SnRK2 regulatory domain packing. **A and B:** Cartoon representation of the ScSAPK10 structure. Highlighted regions represent some of the key regions for kinase activity and/or regulation: ATP binding loop (red), αC (purple), activation loop (orange) and SnRK2-box (green). The hinge region that connects the N- and C-terminal lobes of the kinase domain is colored in yellow. Residues Y165-T181 and D289-M304 were not resolved in the electron density. **C:** Cartoon representation of ScSAPK10 ATP-binding site. The ATP-binding loop (red), the activation loop residues D162(pink) and F163 (orange), as well as the residues K52 (grey) and E67 (purple) related to phosphate transfer, are highlighted. **D:** Structural alignment of ScSAPK10 (gray), Arabidopsis SnRK2.3 (PDB ID: 3UC3 - dark blue) and SnRK2.6 (PDB ID: 3ZUT - light blue).

Superposition of ScSAPK10 onto the structures of AtSnRK2.3 and 2.6 (Ng et al., 2011; Yunta et al., 2011) confirmed our expectation that ABA-responsive SnRK2s from mono and dicot plants are structurally similar - ScSAPK10 and AtSnRK2.3 root mean square deviation (r.m.s.d.): 2.32 Å; ScSAPK10 and AtSnRK2.6 r.m.s.d: 1.35 Å - (Figure 2D). In the crystal, ScSAPK10 adopted an inactive conformation in which the side chain of Phe153 within the conserved kinase motif DFG points towards the ATP-binding site. In this inactive conformation, structurally conserved regions of the kinase domain important for phosphate transfer are kept apart. Moreover, the protein P-loop was found folded towards the kinase hinge region, an orientation that is likely to prevent binding of ATP (Figure 2C).

### SnRK2s-box structure and function are conserved between ScSAPK and AtSnRKs

As seen for other SnRK2 family members, ScSAPK10 SnRK2-box is packed against the αC helix, within the protein kinase domain (Figure 2). Contacts between αC and the SnRK2-box are facilitated by conserved amino acids bearing aliphatic side chains (Figure 3A-C). Previous studies have shown that single-point mutations disturbing hydrophobic interactions between αC and SnRK2-box decreased kinase activity of AtSnRK2s (Ng et al., 2011). Considering the high levels of conservation between ScSAPKs and their *A. thaliana* counterparts, we decided to investigate if a similar mechanism could regulate the activity of the sugarcane proteins.

**Figure 3:**
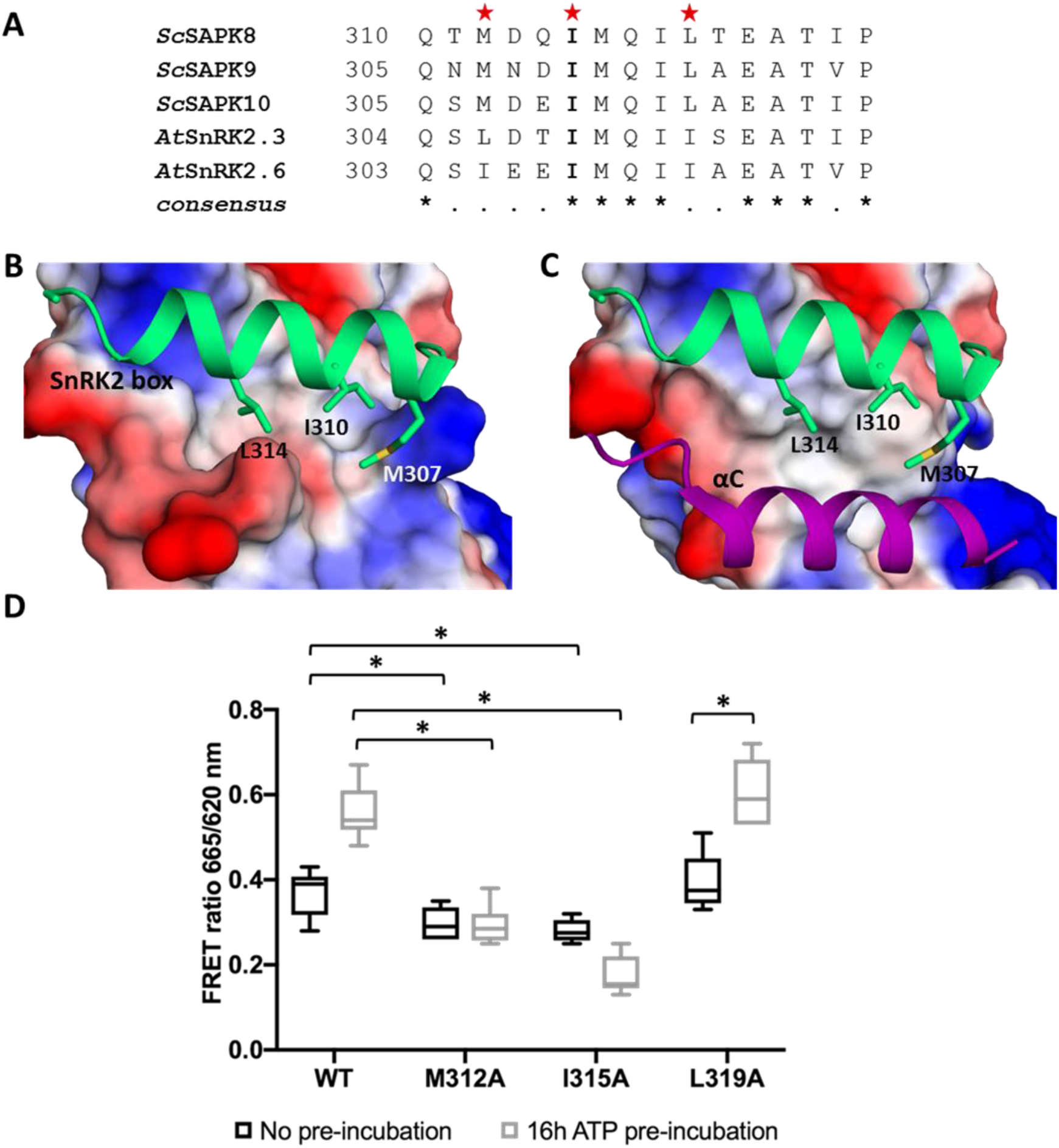
Key residues for SnRK2-box and αC helix interaction are conserved in ScSAPK10 and affect protein activity. **A:** Alignment of SnRK2-box residues from sugarcane SAPKs and SnRK2.6. Red stars represent the residues chosen for site-directed mutagenesis. The residue I315 is conserved in sugarcane and Arabidopsis SnRK2s while the residues M312 and Leu319 are conservatively substituted. **B:** Cartoon representation of SnRK2-box (green) from ScSAPK*10*_Δ*nterm*-Δ*ABA-box* structure. ScSAPK10 SnRK2-box residues M307, I310 and L314 (homologous to ScSAPK8 M312, I315, and L319) are displayed as sticks and make close contact with the αC helix surface. The electrostatic potential analysis shows the negative potential (in red) of the αC surface. The positive potential is represented in blue. **C:** Cartoon representation of SnRK2-box (green) and the αC helix (purple) from ScSAPK*10*_Δ*nterm*-Δ*ABA-box* structure. The electrostatic potential of αC surface was hidden to show the helix position. **D:** Box plot of the enzymatic activity of ScSAPK8 WT and the mutants M312A, I315A and L319A after ATP incubation. The data show the quantity of phosphorylated peptide produced, measured by the ratio of fluorescence intensity at 665 nm (streptavidin-XL665 emission excited by phospho-specific Eu-cryptate conjugated antibody) and 620 nm (Eu-cryptate emission). In both assay conditions, the observed activity for ScSAPK8 WT was significantly higher than the mutants M312A (\**p* = 0.0416 for no ATP pre-incubation and \**p* < 0.0001 for 16h hours ATP pre-incubation) and I315A (\**p* = 0.0114 for no ATP pre-incubation and \**p* < 0.0001 for 16h hours ATP pre-incubation). The L319A activity was similar to WT in both conditions but significantly increased with 16 hours of ATP pre-incubation (\**p* < 0.0001).

To measure the activity of recombinantly expressed ScSAPKs we employed a commercially-available enzymatic assay (KinEASE, Cisbio) and a generic peptide substrate (STK1 from the same vendor). Amongst the three sugarcane proteins, ScSAPK8 was the most active enzyme in this assay (data not shown). We thus decided to study the impact of disrupting the interaction between SnRK2-box and αC helix on the activity of ScSAPK8.

We used site-directed mutagenesis to substitute conserved SnRK2-box residues (Met312, Ile315 or Leu319) with an alanine. Under the experimental condition with no pre-incubation with ATP, the activities of ScSAPK8 mutants M312A and I315A were statistically lower than the observed to wild type enzyme (one-way ANOVA *post-hoc* Dunnett’s, n = 6, *p* = 0.0416 comparing WT to M312A and *p* = 0.0114 for WT and I315A; ANOVA *p* = 0.0011) whereas L319A activity was comparable to wild type (*p* = 0.7021) (Figure 3D). AtSnRK2s are known to be activated by autophosphorylation (Fujii et al., 2009; Ng et al., 2011). We then investigated the impact of pre-incubating wild-type and mutant ScSAPK8s with ATP (16 hours at 25°C) before assaying their activity. Pre-incubation with ATP statistically increased the baseline activity of wild-type and the L319A mutant (two-way ANOVA *post-hoc* Bonferroni’s, *p* < 0.0001, n = 6), whereas activity for mutant M312A remained unaltered (*p* > 0.9999). Surprisingly, the overall activity of I315A was significantly reduced (*p* = 0.0034) (Figure 3D).

To verify if the reduced activity observed for mutants M312 and I315A resulted from the inability of these proteins to self-activate, we used LC-MS to obtain the intact masses of wild-type and mutant proteins. The total number of phosphosites observed after 16-hour incubation with ATP remained the same for wild-type and mutant proteins M312A and I315A (Supplementary Figure S3, S4, and S5). Thus, the reduced activities observed for these two mutant proteins were not due to a defect in their auto-phosphorylation abilities. Surprisingly, mutant L319A displayed three additional phosphosites compared to the wild-type and the other two mutants investigated (Table 2; Supplementary Figure S3 and S6).

**Table 2:**
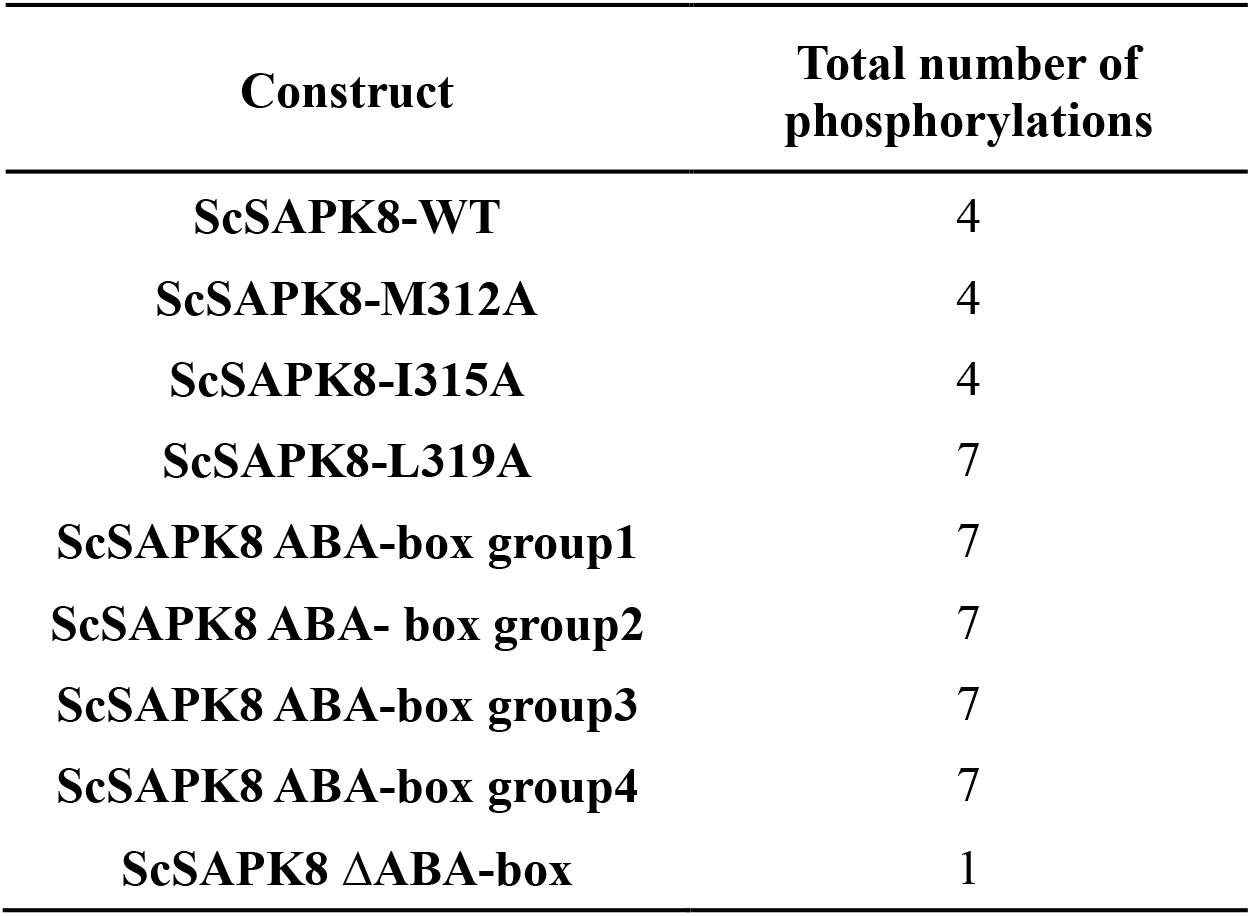
Intact mass analysis of ScSAPK8 proteins after overnight incubation with Mg^2+^/ATP

Taken together, our results indicate that mutations designed to disrupt the interaction between the αC helix and SnRK2s-box coordination in sugarcane SAPK8 did not abolish enzyme activity. Nevertheless, changes in residues Met312 and Ile315 did reduce the overall protein activity after a longer (16-hour) period in the presence of ATP, an effect that might be due to the mutant proteins having lower stability than the wild-type or the L319A mutant.

### Deletion of SAPK8 ABA-box does not directly affect its activity

In addition to the SnRK2-box, another conserved C-terminal region is involved in SnRK2 regulation in dicot plants – the ABA-box (Belin et al., 2006; Boudsocq et al., 2007; Soon et al., 2012). This region mediates the interaction between SnRK2s and PP2C phosphatases, leading to kinase inactivation via dephosphorylation of an essential activation loop serine residue and preventing substrate access to the kinase catalytic site (Belin et al., 2006; Soon et al., 2012). Here we investigated ScSAPK8 ABA-box contribution to kinase activity in the absence of a PP2C phosphatase. For that, we used the enzymatic assay described above to assess the activity of several ScSAPK8 C-terminal mutants designed to either completely remove the protein ABA-box or disrupt the region’s acidic character via replacement of conserved acidic residues with alanines (Figure 4A).

**Figure 4:**
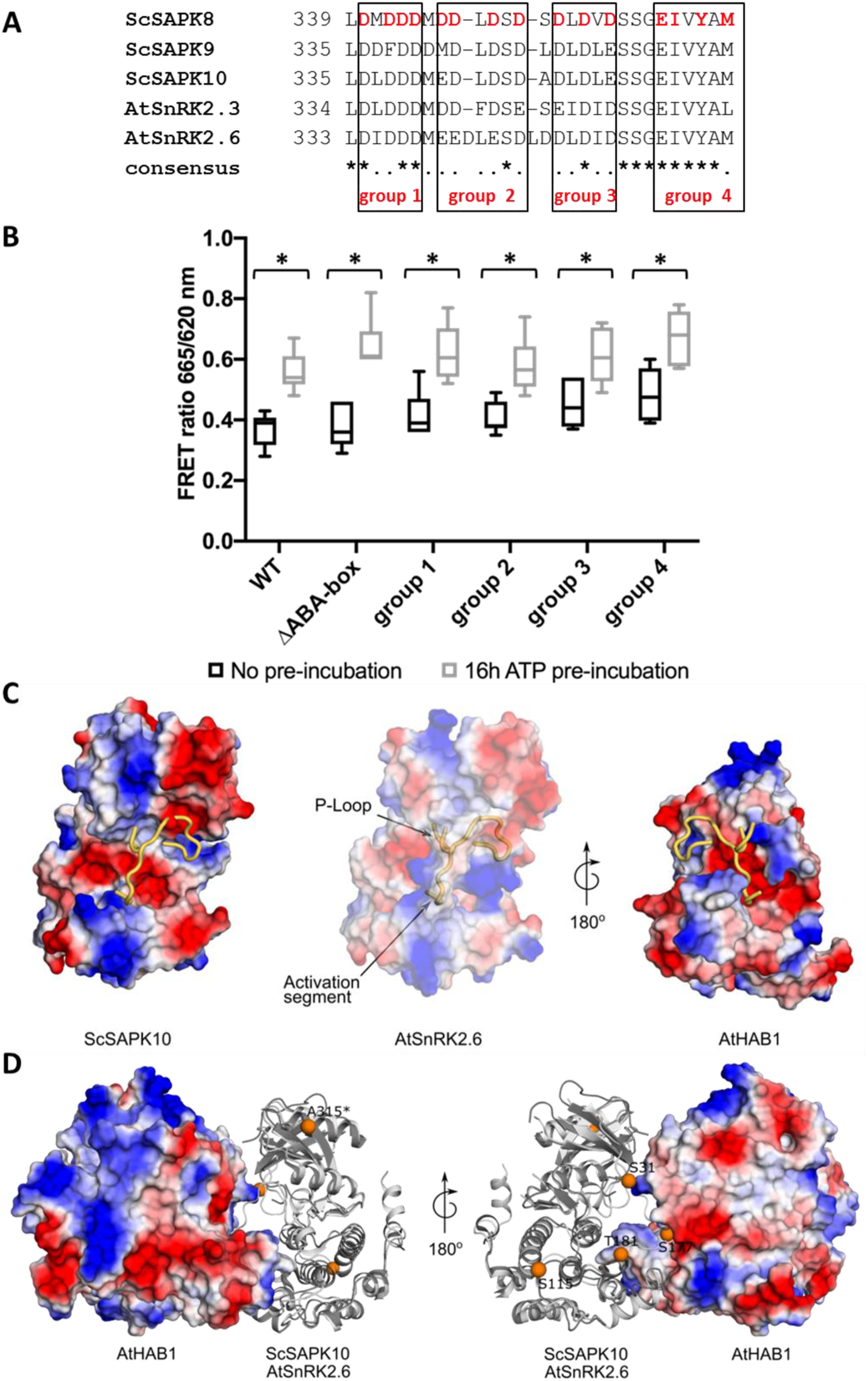
ScSAPK8 ABA-box mutations do not affect protein activity and might affect kinase interaction with PP2C phosphatase. **A:** Alignment of ABA-box residues from sugarcane SAPKs and SnRK2.6. Mutations performed in ScSAPK8 ABA-box are displayed in red and were distributed in four different groups, named group 1 to group 4. In the mutants from group 1 to group 3 all aspartic acid residues (D, in red) were replaced by alanine residues. In group 4, the residues of glutamic acid, isoleucine, tyrosine and methionine (respectively, E, I, Y and M, in red) were mutated to alanine **B:** Box plot of ScSAPK8 WT and ABA-box mutant enzymatic activity after ATP incubation. The data show the quantity of phosphorylated peptide produced, measured by the ratio of fluorescence intensity at 665 nm (streptavidin-XL665 emission excited by phospho-specific Eu-cryptate conjugated antibody) and 620 nm (Eu-cryptate emission). The analysis shows no statistically significant difference between the activity of WT and all the mutants tested. All the proteins presented significantly increased activity after 16 hours of ATP pre-incubation compared to the condition with no pre-incubation (*p* < 0.0001). **C:** Cartoon representation of ScSAPK10, AtSnRK2.6 and AtHAB1 protein surfaces. The electrostatic potential analysis shows the positive potential (in blue) of the protein surfaces around the activation segment and P-loop. **D:** Cartoon representation of ScSAPK10 (dark gray) aligned with AtSnRK2.6 (light gray) and AtHAB1 (represented as electrostatic surface). The orange spheres represents, in the ScSAPK10 structure, the homologous phosphosites identified to ScSAPK8 by mass spectrometry. The ScSAPK10 residues S31, S115, S177, T181 and A315 correspond to S36, S120, S182, T186 and T320 in ScSAPK8 sequence, respectively.

All ScSAPK8 mutants displayed similar overall activity to the wild-type protein. Likewise, pre-treatment with ATP (16 hours at 25 °C) significantly increased enzyme activity for all proteins tested (two-way ANOVA *post-hoc* Bonferroni’s, *p* < 0.0001, n = 6) (Figure 4B). We also used LC-MS to assess the impact of ScSAPK8 ABA-box on protein auto-phosphorylation. Interestingly, more phosphosites could be detected for ScSAPK8 point mutants than for the wild-type protein (7 versus 4 phosphosites, respectively) (Table 2; Supplementary Figure S3, S8 to S11). On the other hand, a single phosphorylation was detected in the truncated version of ScSAPK8 completely lacking the ABA-box in the intact mass analysis (Table 2 and Supplementary Figure S7). Similar results were observed for the ScSAPK10 ΔABA-box protein (data not shown).

Taken together, the data above suggest that ScSAPK8 ABA-box is important for protein overall phosphorylation state, but, by itself, this conserved motif does not regulate enzyme activity.

### Structure of SAPK10 suggests a conserved interaction mechanism with PP2C-type phosphatases

Structural studies revealed that AtSnRK2.6 and PP2C-type phosphatases display complementary electrostatic surfaces at the complex interface (Soon et al., 2012). An overlay of the structures of ScSAPK10 and SnRK2.6 bound to a PP2C-type phosphatase (AtHAB1), revealed that both kinases display similar electrostatic surfaces within the SnRK2.6 region known to interact with PP2C-type phopshatases (Figure 4C).

We used LC-MS/MS to identify sites of autophosphorylation within ScSAPK8 WT and ΔABA-box mutant (Table 3) and then mapped these onto our ScSAPK10 crystal structure. Three (Ser36, Ser182, and Thr186) out of the 5 identified phosphosites in ScSAPK8 WT have structural equivalents in the Arabidopsis protein that are within the kinase:phosphatase complex interface (Figure 4D) (Soon et al., 2012). Curiously, only two phosphosites were observed in ScSAPK8 ΔABA-box mutant, both located in the kinase:phosphatase complex interface.

**Table 3:**
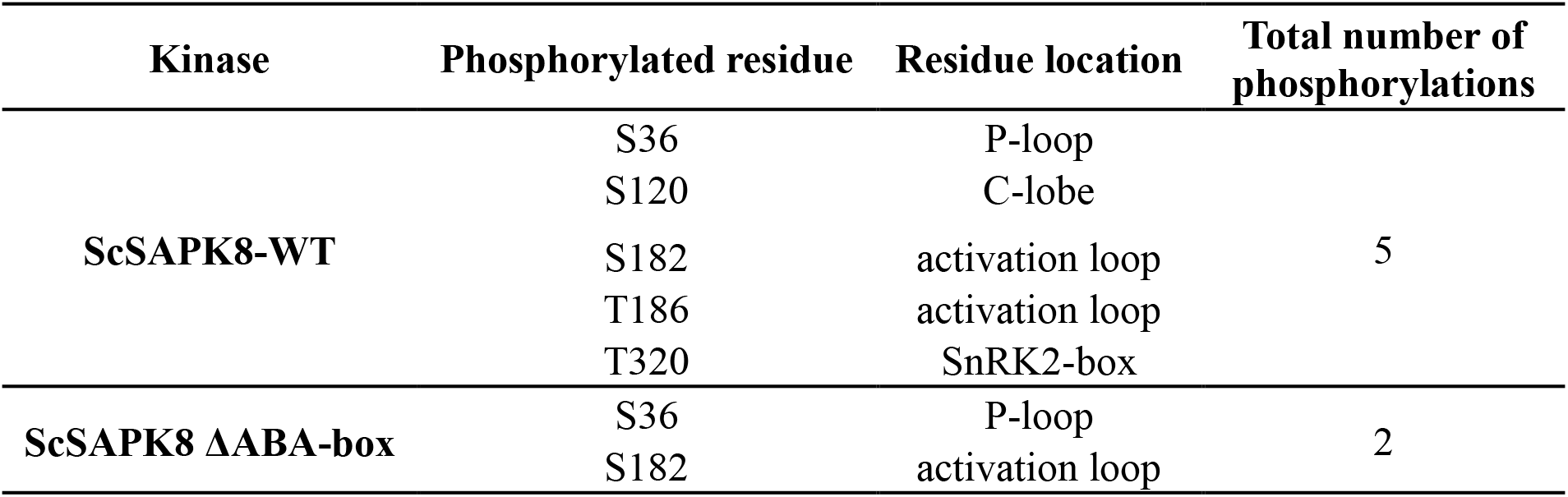
ScSAPK8 phosphopeptides identification by mass spectrometry

| Kinase | Phosphorylated residue | Residue location | Total number of phosphorylations |
| --- | --- | --- | --- |
| ScSAPK8-WT | S36 | P-loop | 5 |
|  | S120 | C-lobe |  |
|  | S182 | activation loop |  |
|  | T186 | activation loop |  |
|  | T320 | SnRK2-box |  |
| ScSAPK8 ΔABA-box | S36 | P-loop | 2 |
|  | S182 | activation loop |  |

These analyses suggest that the overall mechanism regulating the interaction of ABA-responsive kinases and PP2C-type phosphatases are conserved between Arabidopsis and sugarcane proteins.

## DISCUSSION

ABA is a key hormone in both mono- and dicotyledon plants. In dicotyledons, members of the SnRK2 family of protein kinases play a central role in ABA signaling and act as positive regulators of this stress hormone (Fujii et al., 2009; Fujii and Zhu, 2009; Fujita et al., 2009; Nakashima et al., 2009). It is expected that the signaling pathway relaying ABA stimuli is also conserved in monocot plants. Supporting this hypothesis, recent studies have shown that ABA strongly activates expression of SnRK2 counterparts in sugarcane (Li et al., 2017). Here we confirmed that three functional SnRK2 proteins - ScSAPK8, ScSAPK9, and ScSAPK10-are encoded by the sugarcane genome; further suggesting that the ABA-response pathway is conserved in both mono- and dicotyledons.

Our analyses of the ScSAPK structure and biochemistry strongly suggest that ABA-responsive kinases in sugarcane are functionally equivalent to their counterparts in Arabidopsis. The structure of ScSAPK10 revealed that the C-terminal SnRK2-box folds into an α-helix and interacts with the structurally-conserved αC from the protein kinase domain. A similar interaction has been reported for AtSnRK2 proteins and is thought important for kinase activity, akin to the activation mechanism of human cyclin-dependent kinases (Jeffrey et al., 1995; Ng et al., 2011). Nevertheless, the ScSAPK structure determined here and previously determined SnRK2 structures have captured the protein in its inactive kinase state, despite the observed interaction between αC and SnRK2-box. Obtaining the structure of an SnRK2 family member in an active conformation would shed light on how SnRK2-box contributes to protein activity.

Biochemical assays have shown that disrupting hydrophobic contacts between αC and SnRK2-box via point mutations in AtSnRK2 abolished protein activity (Belin et al., 2006; Ng et al., 2011; Soon et al., 2012; Yunta et al., 2011). However, equivalent mutations in ScSAPK8 SnRK2-box had no to little effect on protein activity following no pre-incubation with ATP. Although we did see reduced activity for some of the SnRK2-box mutants after a 16-hour pre-incubation period with ATP, we cannot discard the possibility this reduced activity was due to differences in stability between wild-type and mutant proteins. Nevertheless, our results suggest that either the introduced mutations did not disrupt the SnRK2-box αC interaction in ScSAPK8 or that this region is not critical for protein activity. It is difficult to conclude from *in vitro* experiments performed using different assays, and we believe this issue requires further exploration. More importantly, how, and if, the SnRK2-box interaction to αC is modulated *in vivo* remains to be elucidated.

A second region important for regulating SnRK2 activity is the ABA-box. This region is a stretch of mostly acidic residues that mediate the direct interaction between SnRK2 proteins and basic patches on the surface of PP2C-type phosphatases (Soon et al., 2012). This interaction prevents the phosphorylation activity of SnRK2s. *In vitro*, total deletion of AtSnRK2.6 ABA-box did not affect protein activity. However, ectopic expression of this truncated protein in Arabidopsis *snrk2.6* mutants could not restore stomatal closure response. These same studies revealed that phosphorylation of sites within the kinase domain was important for promoting wild-type response to ABA (Belin et al., 2006; Yoshida et al., 2006).

Our data indicated that deleting ABA-box from ScSAPK8 did indeed reduce the overall number of autophosphorylation sites within this protein - from 5 in the wild-type protein to two in the ΔABA-box mutant. Nevertheless, deletion of ScSAPK8 ABA-box did not alter kinase activity on a generic peptide substrate, suggesting that autophosphorylation of residues located in the P-loop (Ser36) and activation loop (Ser182) might be sufficient for full kinase activity *in vitro*. However, the lack of activity of the ΔABA-box SnRK2 mutant in Arabidopsis might indicate a role of the additional phosphorylation events in kinase activity *in vivo*. In addition, the peptide analysis (Table 3) of ScSAPK8 WT and ΔABA-box mutant presented an increased number of phosphorylation sites for both proteins when compared to intact mass analysis (Table 2). The intact mass analysis represents the measurement of all different phospho-states of a certain protein in a mixture while the peptide analysis allows the precise identification of phosphosite position. In our work, the discrepancy between the intact mass and the number of phosphosites identified for ScSAPK8 WT and ΔABA-box mutant might indicate that S36 and S182 phosphorylation could not co-exist in the same molecule of protein. In this scenario, the intact mass analysis would result in a single phospho-state corresponding to two different phosphosites only unveil after protein digestion. The underlying mechanism behind these observations is not clear at this moment and will require further investigation.

The activation loop is a structurally conserved feature of protein kinases, and phosphorylation of key residues within this region stabilizes the protein in an active conformation (Nolen et al., 2004). In Arabidopsis SnRK2s, phosphorylation of a serine residue (Ser175 in SnRK2.6) within the protein activation loop is essential for kinase activation (Belin et al., 2006; Boudsocq et al., 2007; Ng et al., 2011; Vlad et al., 2009).

We identified the equivalent residue in ScSAPK8 as phosphorylated after incubation with Mg^2+^/ATP, further suggesting that SnRK2 family members from both monocots and dicots display similar regulatory mechanisms.

## CONCLUSION

Here, we determined the crystallographic structure and performed the biochemical characterization of ABA-related SnRK2 proteins from the crop plant sugarcane. Our analyses suggest a role for ScSAPK ABA-box in protein autophosphorylation but not in overall enzyme activity. Moreover, disrupting the SnRK2-box:αC interaction had no to little effect in ScSAPK8 kinase activity, despite the structural conservation between sugarcane and Arabidopsis proteins. Future studies are required to evaluate the role of these two conserved regions, as well as that of the multiple phosphosites identified here, in kinase activation and activity *in planta*.

## Supporting information

Supplementary material_Righetto

## Author contributions

Germanna Righetto participated in all parts of the project. Dev Sriranganadane performed all mass spec analysis. Levon Halabelian collected diffraction data. Carla G. Chiodi helped with protein expression and purification. Opher Gileadi coordinated the design of expression constructs. Rafael M. Counãgo coordinated crystal structure determination and refinement. Katlin B. Massirer, Jonathan Elkins, Marcelo Menossi and Rafael M. Counãgo coordinated the project. Germanna Righetto and Rafael M. Counãgo wrote the manuscript. All authors revised the manuscript.

## Acknowledgments

This research used the Pilatus 6M detector on 19-ID beamline of the Advanced Photon Source, a U.S. Department of Energy (DOE) Office of Science User Facility operated for the DOE Office of Science by Argonne National Laboratory under Contract No. DE-AC02-06CH11357. We thank the staff of the Life Sciences Core Facility (LaCTAD) from State University of Campinas (UNICAMP), for the genomics and proteomics analysis.

## Data Availability Statement

The coordinates and structure factors for ScSAPK10 crystal structure reported here have been deposited in the Protein Data Bank with accession code 5WAX.

