## Supplementary material_Righetto for "The C-terminal domains SnRK2-box and ABA-box have a role in sugarcane SnRK2s auto-activation and activity"

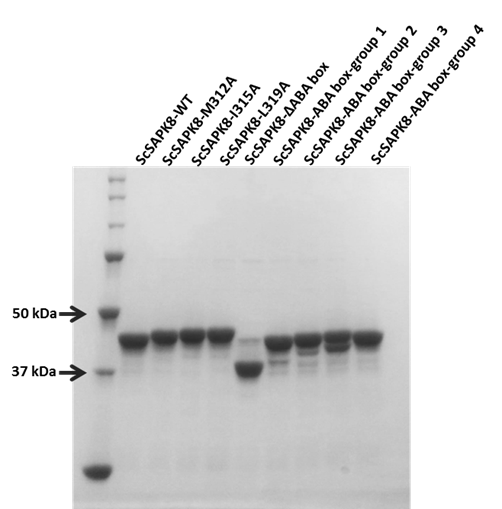

**Supplementary Figure S1: Coomassie stained SDS-PAGE of ScSAPK8 WT and mutants.** The image represents all the proteins after affinity purification and dilution to 20 μM final concentration. The protein concentration was estimated by the Bradford method (Sigma-Aldrich) before gel loading.

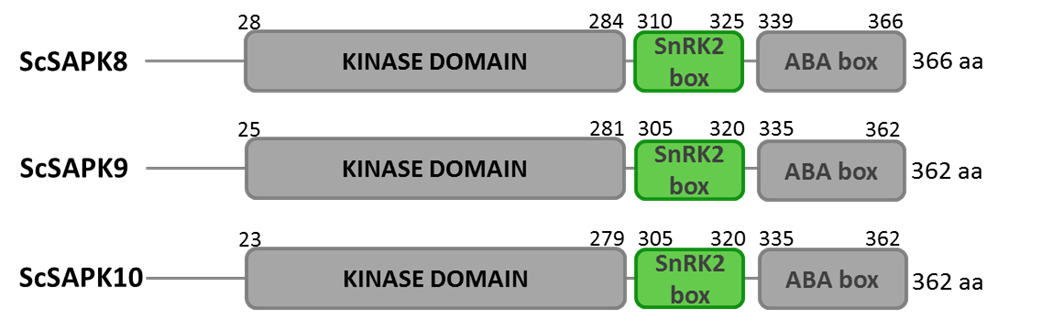

Supplementary Figure S2: Schematic representation of full-length ScSAPK8, ScSAPK9, and ScSAPK10. Each protein has an N-terminal kinase domain and a C-terminal region containing the regulatory domains SnRK2-box and ABA-box. Numbers represent the amino acid positions of each protein domain.

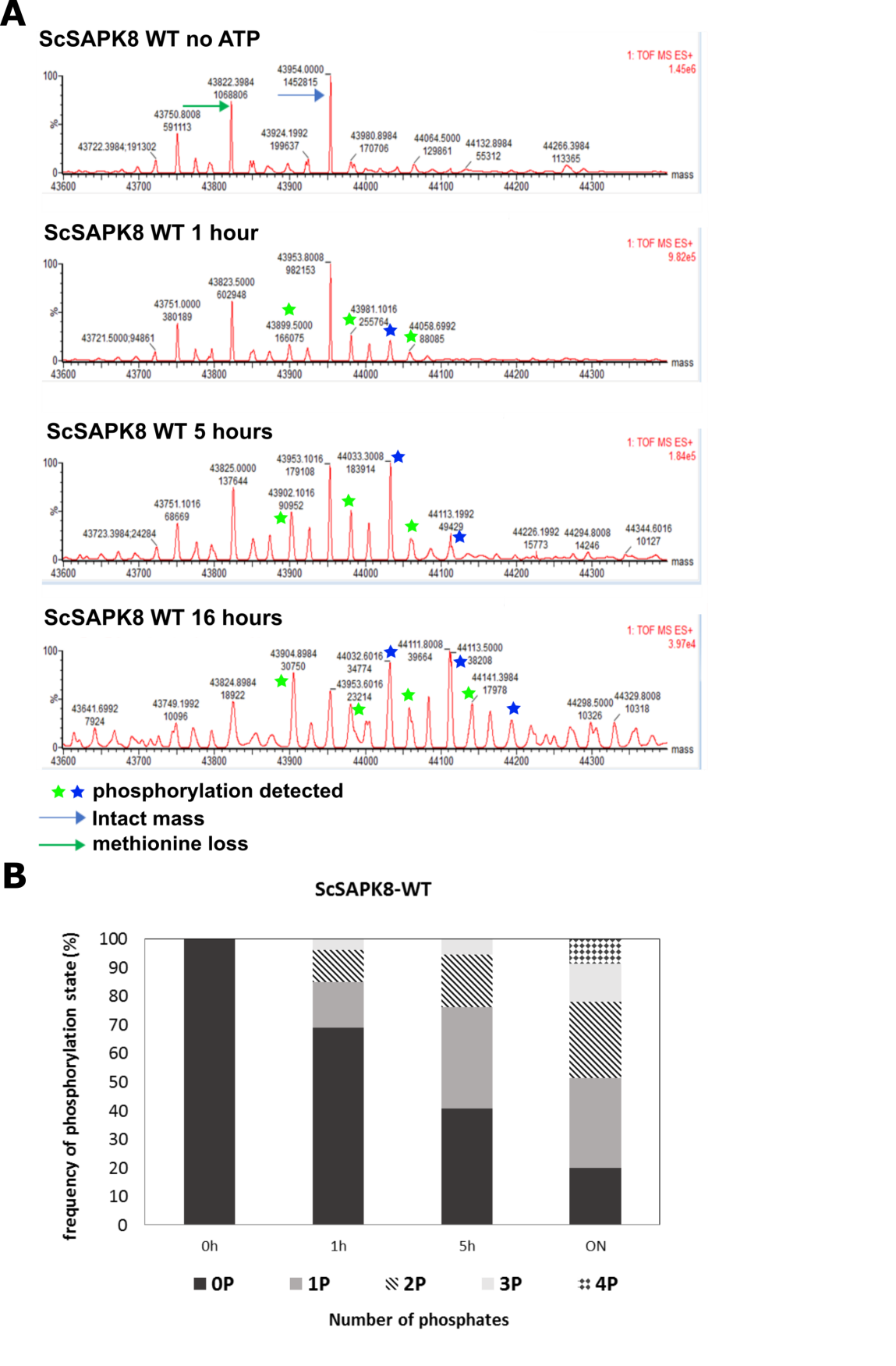

**Supplementary Figure** **S3: ScSAPK8 WT autophosphorylation. A:** Deconvoluted mass spectrum at time 0, 1 hour, 5 hours and overnight, generated by MAX ENT1 software (Waters). The blue arrow represents the intact kinase mass, and the blue stars represent the detected masses with one or more phosphorylations. The green arrows indicate the exact kinase mass with the loss of methionine that could occur during protein expression. The green stars represent masses of this kinase form with one or more phosphorylations. **B:** Graphical representation of protein autophosphorylation over time. The percentage values were calculated using the ion counts extracted from the deconvoluted spectrum.

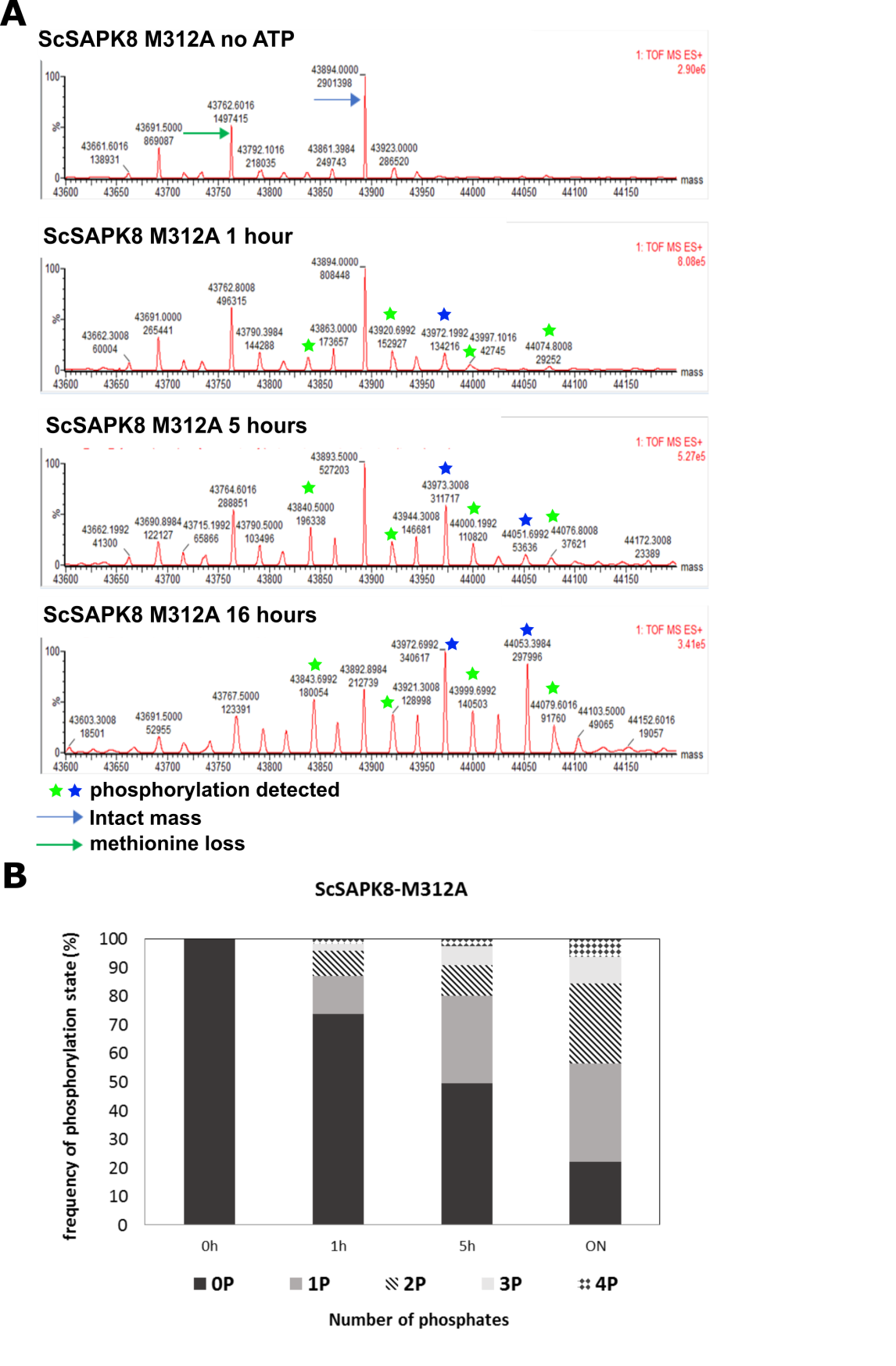

**Supplementary Figure S4: ScSAPK8-M312A autophosphorylation. A:** Deconvoluted mass spectrum at time 0, 1 hour, 5 hours and overnight, generated by MAX ENT1 software (Waters). The blue arrow represents the intact kinase mass, and the blue stars represent the detected masses with one or more phosphorylations. The green arrows indicate the exact kinase mass with the loss of methionine that could occur during protein expression. The green stars represent masses of this kinase form with one or more phosphorylations. **B:** Graphical representation of protein autophosphorylation over time. The percentage values were calculated using the ion counts extracted from the deconvoluted spectrum.

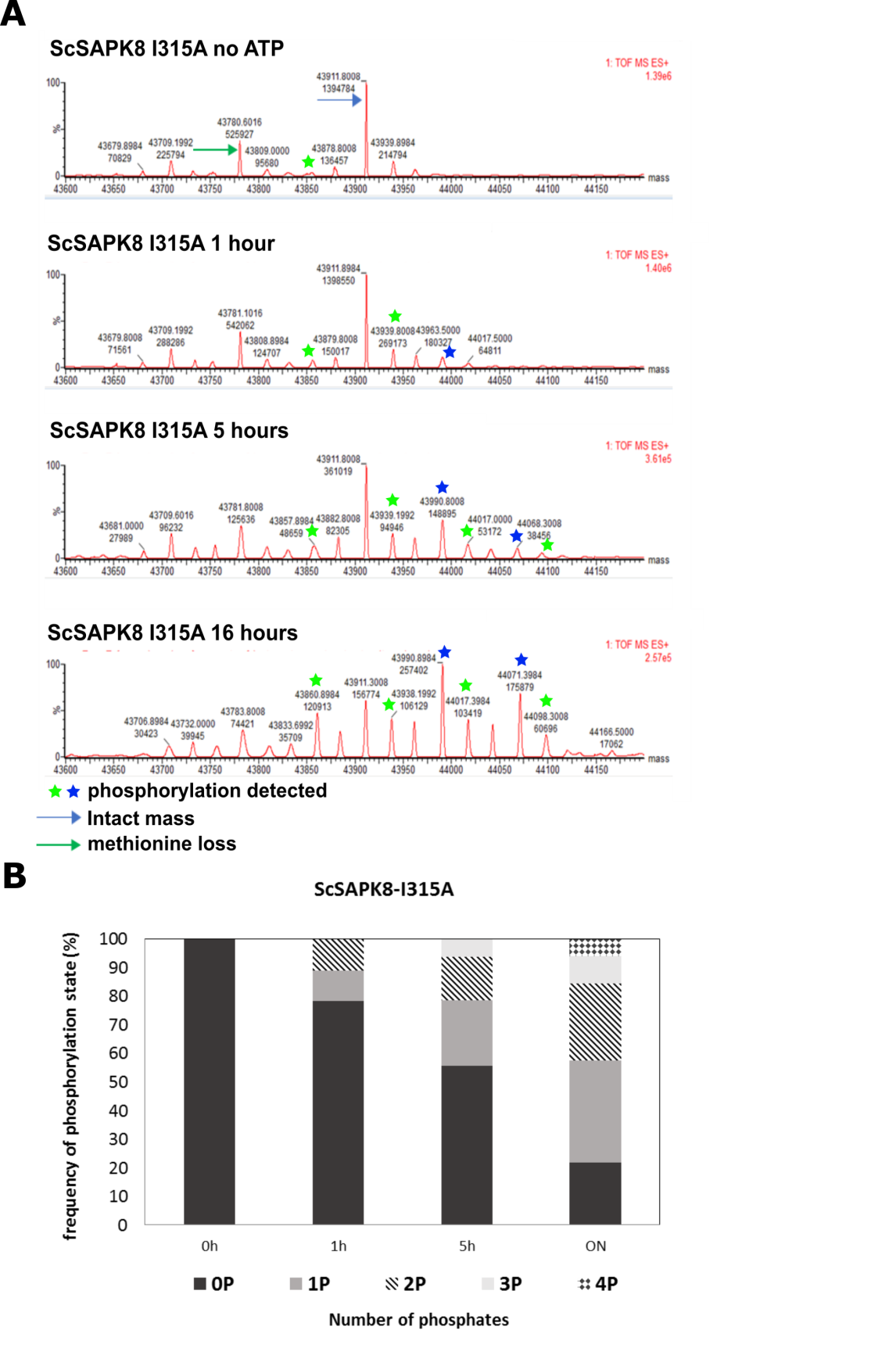

**Supplementary Figure S5: ScSAPK8-I315A autophosphorylation. A:** Deconvoluted mass spectrum at time 0, 1 hour, 5 hours and overnight, generated by MAX ENT1 software (Waters). The blue arrow represents the intact kinase mass, and the blue stars represent the detected masses with one or more phosphorylations. The green arrows indicate the exact kinase mass with the loss of methionine that could occur during protein expression. The green stars represent masses of this kinase form with one or more phosphorylations. **B:** Graphical representation of protein autophosphorylation over time. The percentage values were calculated using the ion counts extracted from the deconvoluted spectrum.

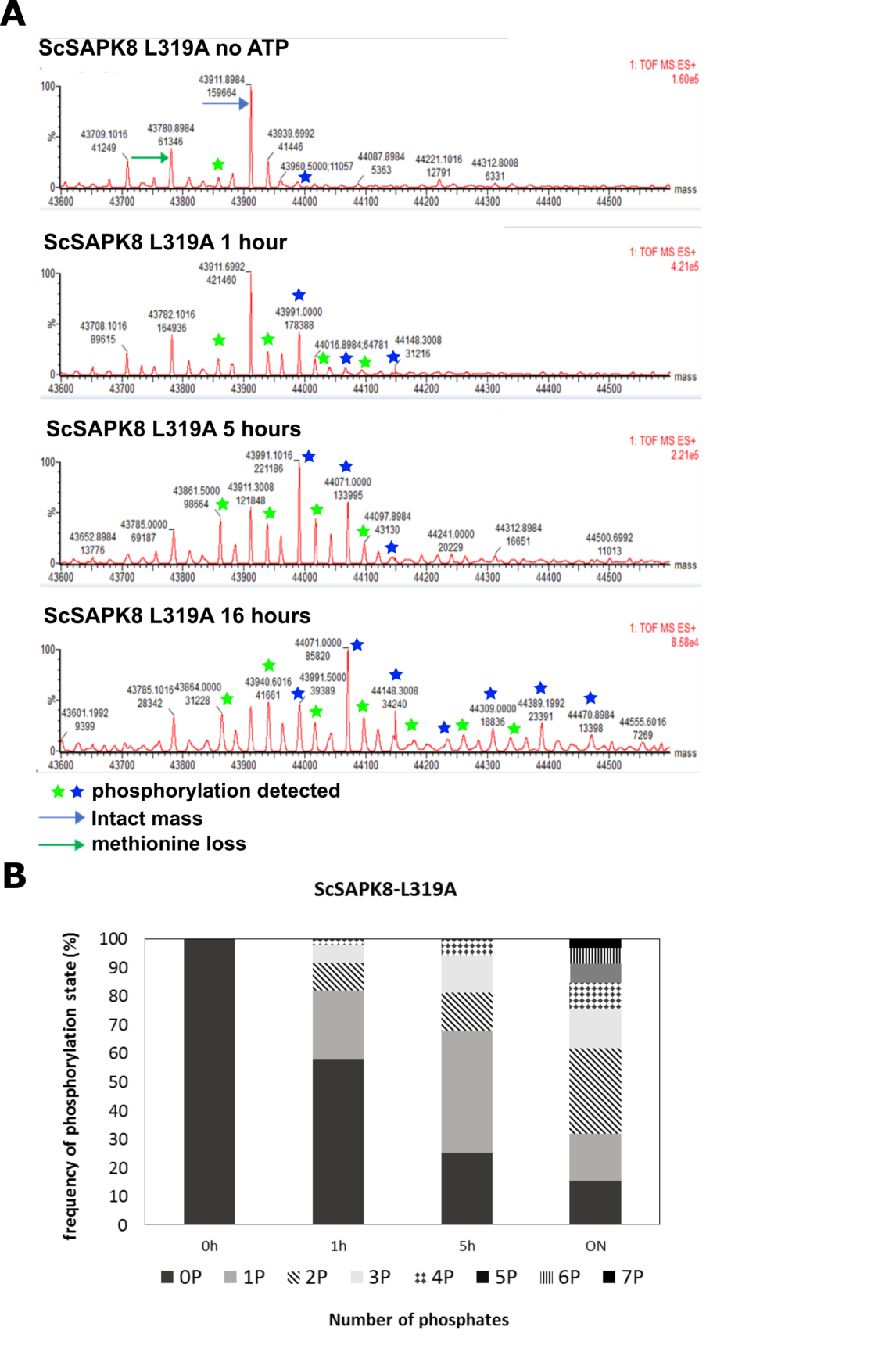

**Supplementary Figure S6: ScSAPK8-L319A autophosphorylation. A:** Deconvoluted mass spectrum at time 0, 1 hour, 5 hours and overnight, generated by MAX ENT1 software (Waters). The blue arrow represents the intact kinase mass, and the blue stars represent the detected masses with one or more phosphorylations. The green arrows indicate the exact kinase mass with the loss of methionine that could occur during protein expression. The green stars represent masses of this kinase form with one or more phosphorylations. **B:** Graphical representation of protein autophosphorylation over time. The percentage values were calculated using the ion counts extracted from the deconvoluted spectrum.

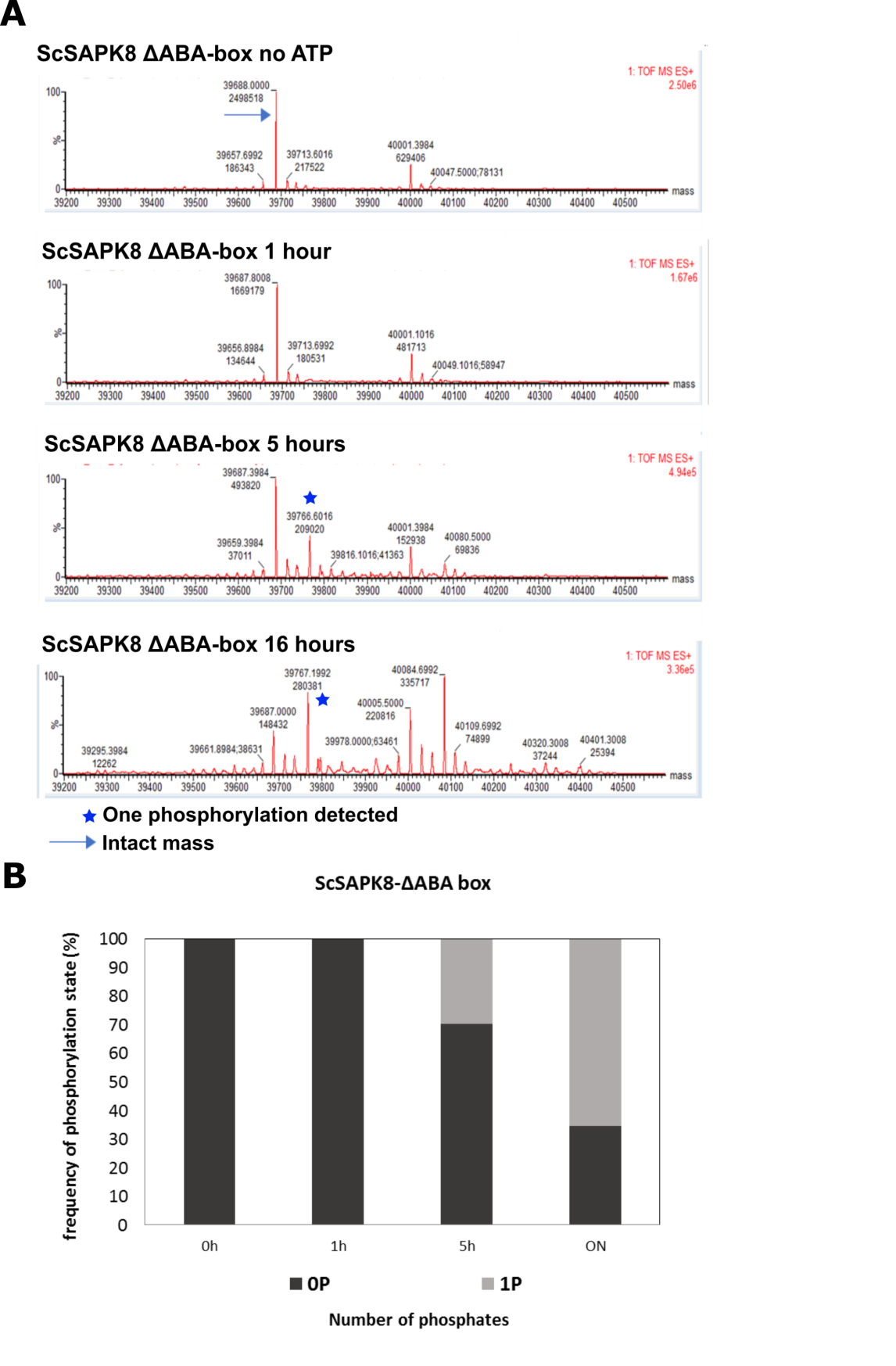

**Supplementary Figure S7: ScSAPK8-∆ABA-box autophosphorylation. A:** Deconvoluted mass spectrum at time 0, 1 hour, 5 hours and overnight, generated by MAX ENT1 software (Waters). The blue arrow represents the intact kinase mass, and the blue stars represent the detected masses with one or more phosphorylations. **B:** Graphical representation of protein autophosphorylation over time. The percentage values were calculated the ion counts extracted from the deconvoluted spectrum.

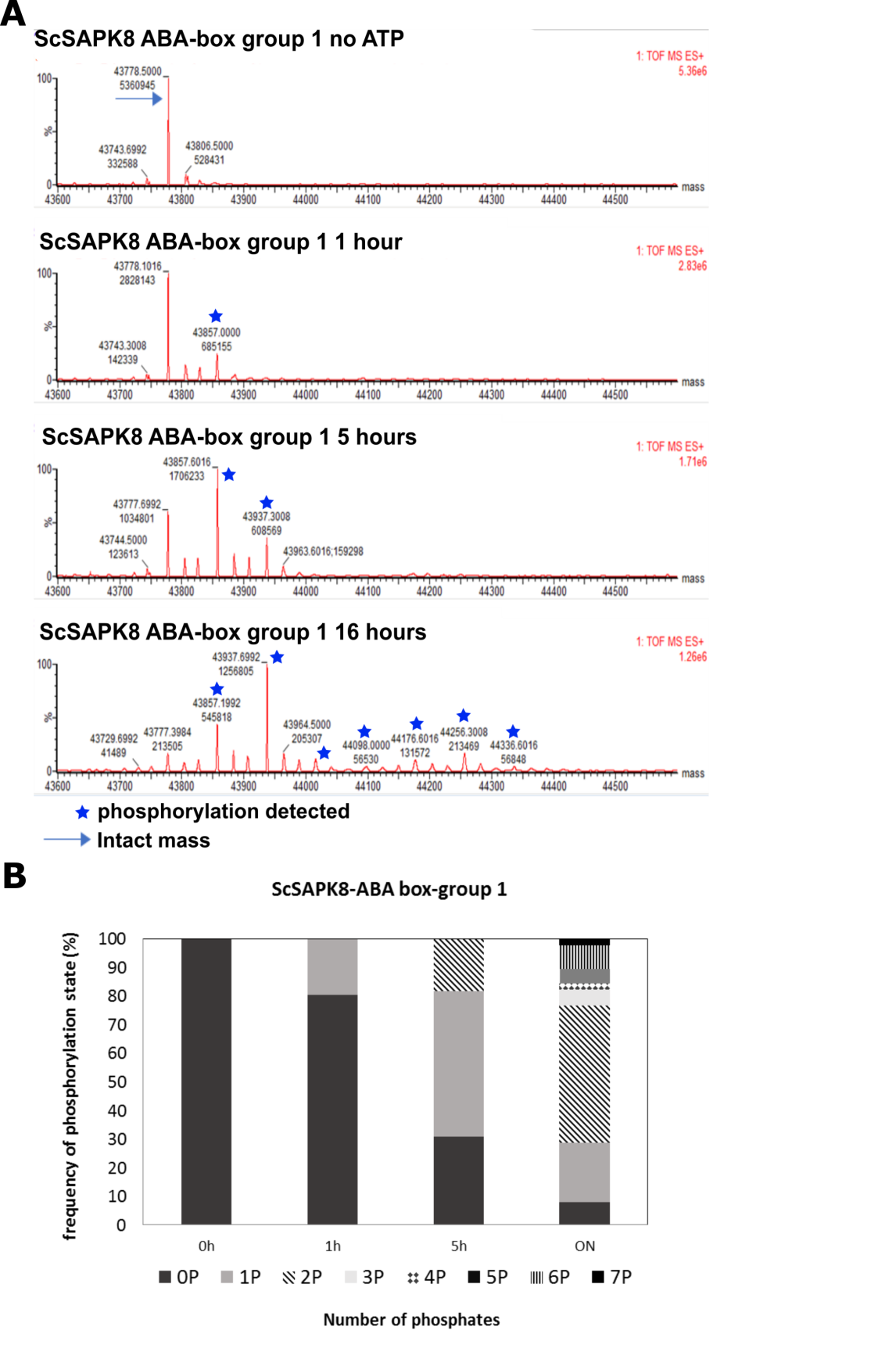

**Supplementary Figure S8: ScSAPK8-ABAbox-group1 autophosphorylation. A:** Deconvoluted mass spectrum at time 0, 1 hour, 5 hours and overnight, generated by MAX ENT1 software (Waters). The blue arrow represents the intact kinase mass, and the blue stars represent the detected masses with one or more phosphorylations. **B:** Graphical representation of protein autophosphorylation over time. The percentage values were calculated the ion counts extracted from the deconvoluted spectrum.

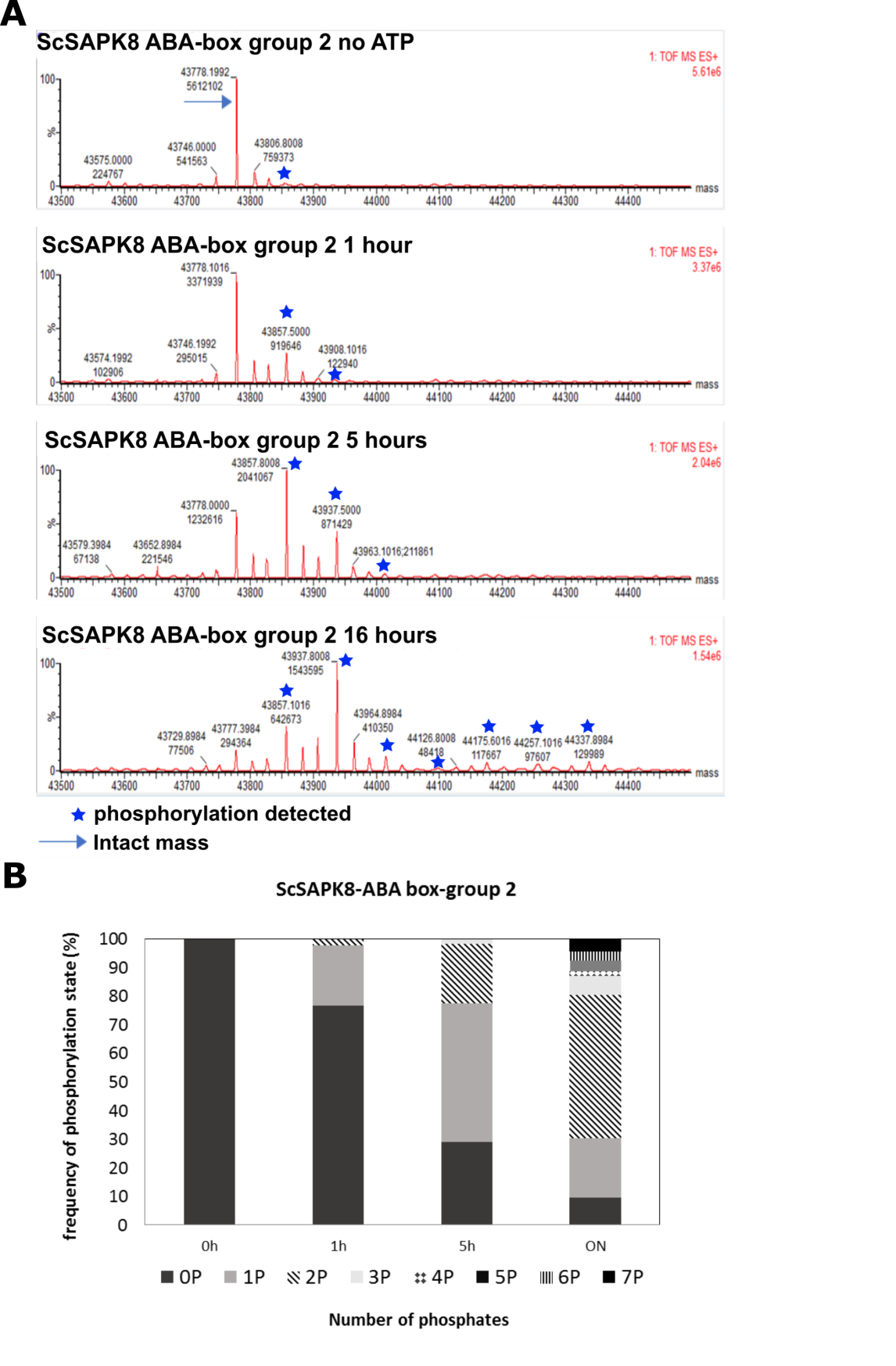

**Supplementary Figure S9: ScSAPK8-ABAbox-group2 autophosphorylation. A:** Deconvoluted mass spectrum at time 0, 1 hour, 5 hours and overnight, generated by MAX ENT1 software (Waters). The blue arrow represents the intact kinase mass, and the blue stars represent the detected masses with one or more phosphorylations. **B:** Graphical representation of protein autophosphorylation over time. The percentage values were calculated the ion counts extracted from the deconvoluted spectrum.

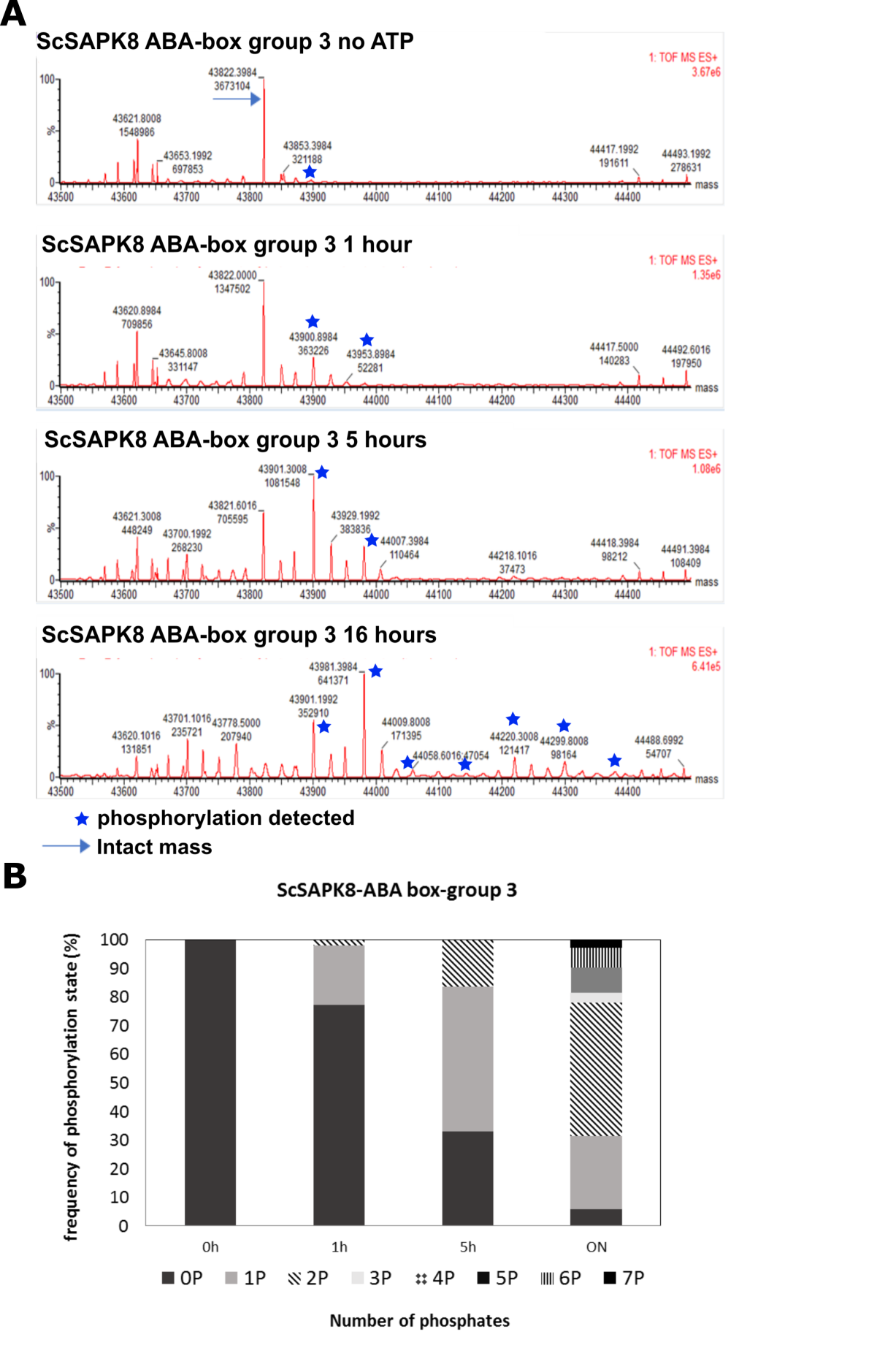

**Supplementary Figure S10: ScSAPK8-ABAbox-group3 autophosphorylation. A:** Deconvoluted mass spectrum at time 0, 1 hour, 5 hours and overnight, generated by MAX ENT1 software (Waters). The blue arrow represents the intact kinase mass, and the blue stars represent the detected masses with one or more phosphorylations. **B:** Graphical representation of protein autophosphorylation over time. The percentage values were calculated the ion counts extracted from the deconvoluted spectrum.

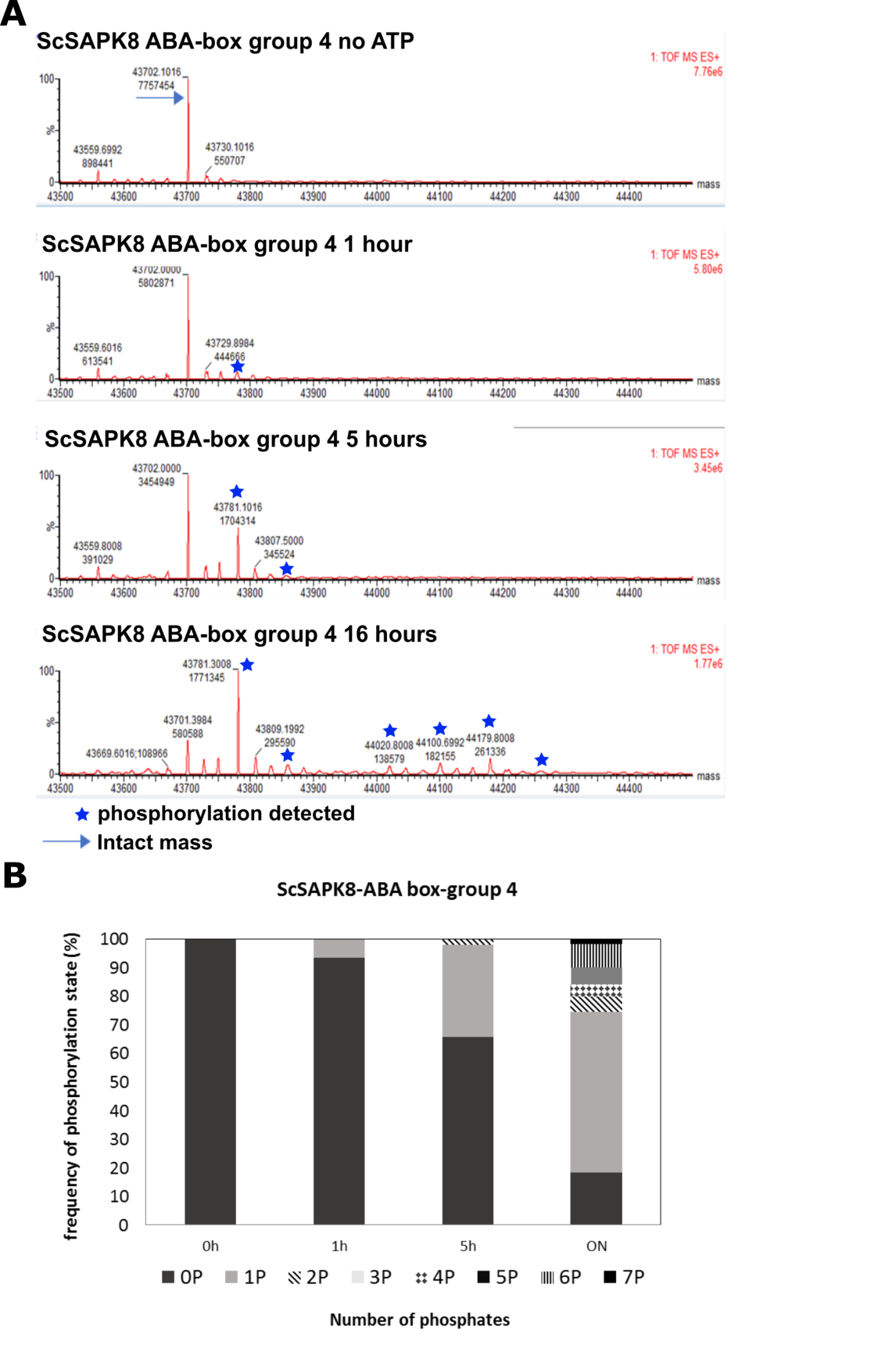

**Supplementary Figure S11: ScSAPK8-ABAbox-group4 autophosphorylation. A:** Deconvoluted mass spectrum at time 0, 1 hour, 5 hours and overnight, generated by MAX ENT1 software (Waters). The blue arrow represents the intact kinase mass, and the blue stars represent the detected masses with one or more phosphorylations. **B:** Graphical representation of protein autophosphorylation over time. The percentage values were calculated the ion counts extracted from the deconvoluted spectrum.

**Table S1: List of primer sequences**

| **Primer Name** | **Sequence 5' - 3'** | **Used for** |
| --- | --- | --- |
| Sc_SAPK8_MluI_NdeI_F | ACGCGTCATATGATGGCAGGGCCGGCGCCG | isolation from cDNA and cloning |
| Sc_SAPK8_NotI_R | GCGGCCGCCATTGCGTACACAATCTCACC |  |
| Sc_SAPK9_MluI_NdeI_F | ACGCGTCATATGATGGCGAGGACGCCGGCA |  |
| Sc_SAPK9_NotI_R | GCGGCCGCCATGGCATACACTATCTCTCC |  |
| Sc_SAPK10_Mlu_NdeI_F | ACGCGTCATATG ATGGACCGGGCGGCGCTC |  |
| Sc_SAPK10_NotI_R | GCGGCCGCCATAGCATACACGATCTCCC |  |
| SAPK8_full_S | TACTTCCAATCCATGGCAGGGCCGGCG | cloning in pNIC28-Bsa4 vector |
| SAPK8_full_AS | TATCCACCTTTACTGTCACATTGCGTACACAATCTCACC |  |
| SAPK9_full_S | TACTTCCAATCCATGGCGAGGACGCCG |  |
| SAPK9_full_AS | TATCCACCTTTACTGTCACATGGCATACACTATCTCTCC |  |
| SAPK10_full_S | TACTTCCAATCCATGGACCGGGCGGCG |  |
| SAPK10_full_AS | TATCCACCTTTACTGTCACATAGCATACACGATCTCCCC |  |
| SAPK10_dm_S | TACTTCCAATCCATGGACATGCCCATAATGCAC | ScSAPK10 truncation |
| SAPK10_dm_AS | TATCCACCTTTACTGTCATGGAATGGTCGCCTCGG |  |
| ScSAPK8_M312A_S | AATGCAGACCGCGGATCAGATCA | SnRK2-box site directed mutagenesis |
| ScSAPK8_M312A_AS | TGATCTGATCCGCGGTCTGCATT |  |
| ScSAPK8_I315A_S | ATGGATCAGGCCATGCAGATTTTG |  |
| ScSAPK8_I315A_AS | CAAAATCTGCATGGCCTGATCCATGG |  |
| ScSAPK8_L319A_S | CATGCAGATTGCGACAGAGGCCA |  |
| ScSAPK8_L319A_AS | TGGCCTCTGTCGCAATCTGCATG |  |
| ScSAPK8_group1_S | GATGGATTGGCCATGGCCGCCGCCATGGATGAT | ABA-box site directed mutagenesis |
| ScSAPK8_group1_AS | ATCATCCATGGCGGCGGCCATGGCCAATCCATC |  |
| ScSAPK8_group2_S | CGACGACATGGCTGCTCTTGCCTCCGCCTCAGATCTTG |  |
| ScSAPK8_group2_AS | CAAGATCTGAGGCGGAGGCAAGAGCAGCCATGTCGTCG |  |
| ScSAPK8_group3_S | TCCGACTCAGCTCTTGCTGTTGCCAGCAGCGGT |  |
| ScSAPK8_group3_AS | ACCGCTGCTGGCAACAGCAAGAGCTGAGTCGGA |  |
| ScSAPK8_group4_S | AGCAGTGGAGCGGCTGTGGCCGCAGCGTGACAGTAAAGGTGGATA |  |
| ScSAPK8_group4_AS | TATCCACCTTTACTGTCACGCTGCGGCCACAGCCGCTCCACTGCT |  |
| SAPK8_full_S | TACTTCCAATCCATGGCAGGGCCGGCG | ABA-box deletion |
| SAPK8_dm_AS | TATCCACCTTTACTGTCAAGGTGGTATGGTGGCCTC |  |
